# Mod(mdg4) variants repress telomeric retrotransposon *HeT-A* by blocking subtelomeric enhancers

**DOI:** 10.1101/2022.01.16.476534

**Authors:** Chikara Takeuchi, Moe Yokoshi, Shu Kondo, Aoi Shibuya, Kuniaki Saito, Takashi Fukaya, Haruhiko Siomi, Yuka W. Iwasaki

## Abstract

Telomeres in *Drosophila* are composed of sequential non-LTR retrotransposons: *HeT-A, TART*, and *TAHRE*. Although they are repressed by the piRNA pathway in the germline, how these retrotransposons are regulated in somatic cells is poorly understood. Here, we show that specific splice variants of Mod(mdg4) repress *HeT-A* by blocking subtelomeric enhancers in ovarian somatic cells. We found that the Mod(mdg4)-N variant represses *HeT-A* expression most efficiently among the variants. Mod(mdg4)-N mutant flies show elevated *HeT-A* expression and female sterility. Mod(mdg4)-N-binding subtelomeric sequences exhibit enhancer-blocking activity, and recruitment of RNA polymerase II (Pol II) on subtelomeres by Mod(mdg4)-N is essential for this enhancer-blocking. Moreover, Mod(mdg4)-N functions to form chromatin boundaries of higher-order chromatin conformation but this mechanism is independent of its Pol II recruitment activity at telomeres/subtelomeres. This study provides a link between enhancer-blocking and telomere regulation, and two different molecular mechanisms exhibited by an insulator protein to orchestrate precise gene expression.

## Introduction

Telomeres are specialized structures that protect the ends of chromosomes, and their loss of function can lead to a variety of defects, including fusion of chromosome ends (De Lange, 2018). In human, telomere sequences are generally composed of short tandem repeats such as TTAGGG and these tandem repeats are elongated by telomerase (Greider and Blackburn, 1985). Telomerase RNA component (TERC) and telomerase reverse transcriptase (TERT) form the telomerase complex (Blasco et al., 1995; Feng et al., 1995; Greider and Blackburn, 1987), which is recruited by telomere binding proteins and TERC is reverse-transcribed at chromosomal ends to add tandem repeats (Jiang et al., 2018). The reverse transcriptase domain has homology with Penelope-like elements (PLE), a class of retroelements (Gladyshev and Arkhipova, 2007; Podlevsky and Chen, 2016), indicating that telomerase might have evolved from retroelements.

In contrast with these telomere maintenance mechanisms, some invertebrate species lack telomerase activity and telomeres are replaced by retrotransposons: *TRAS* and *SART* in *Bombyx mori* (Okazaki et al., 1995; Osanai-Futahashi and Fujiwara, 2011), and *HeT-A, TART* and *TAHRE* in *Drosophila melanogaster* (Abad et al., 2004; Biessmann et al., 1990; Levis et al., 1993; Pardue and Debaryshe, 2002). Telomeres of *D. melanogaster* lack tandem repeat sequences completely and it is proposed that telomere length is maintained by transpositions of these retrotransposons (Kordyukova et al., 2018; Pardue and Debaryshe, 2002; Savitsky et al., 2006) (Fig. 1A). *HeT-A, TART* and *TAHRE* have notable features: (1) they only exist at telomeres, (2) they line up unidirectionally, and (3) *HeT-A* is the most abundant among the three telomeric retrotransposons.

**Fig. 1.**
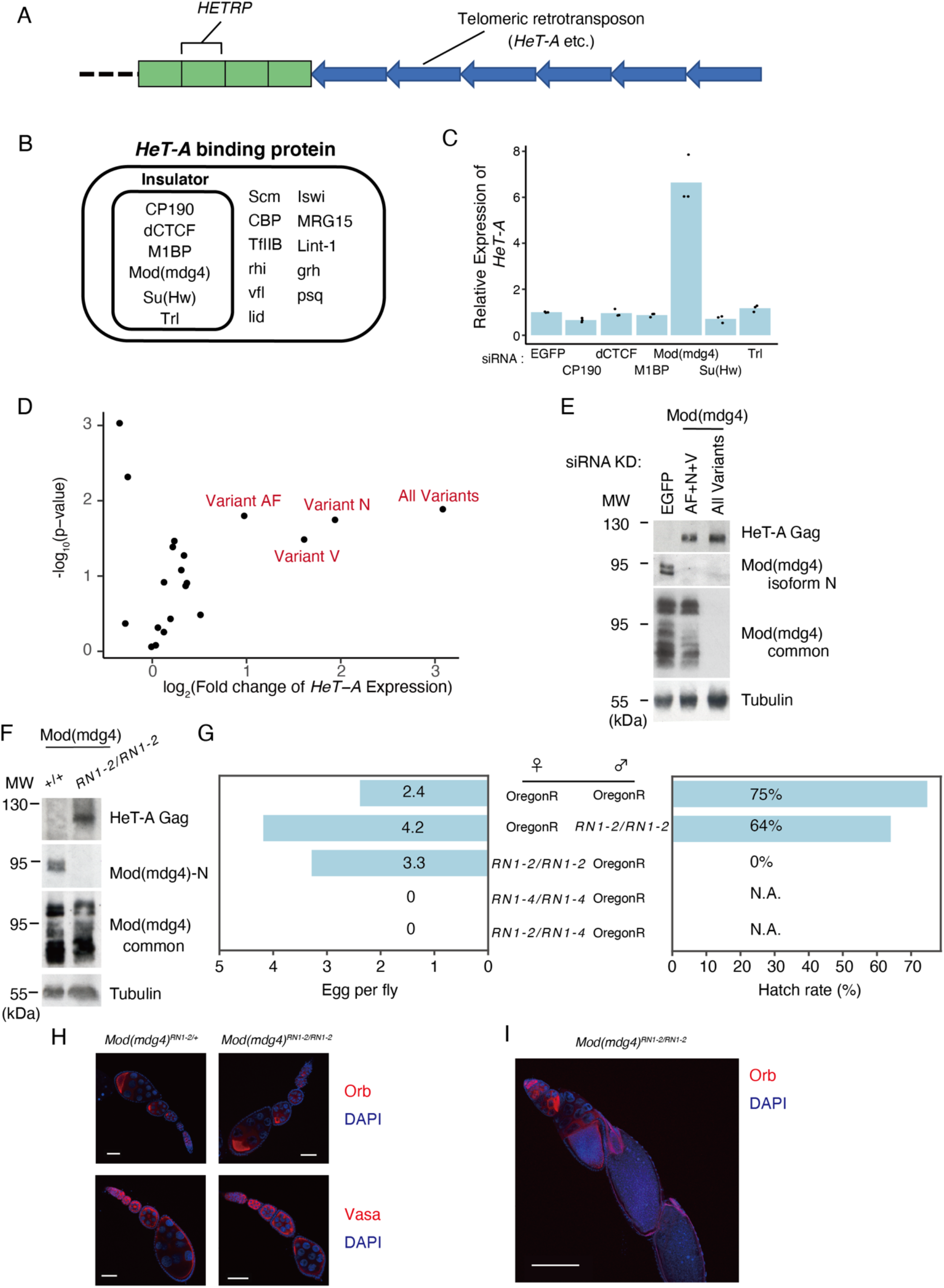
Mod(mdg4) restricts *HeT-A* expression in a variant-specific manner in both OSC and ovaries. **A**, Schematic representation of typical *D. melanogaster* telomere/subtelomere structure. Especially on the right arms of chromosomes 2 and 3, subtelomeres contain several *HETRP* sequences. Next to these subtelomeric *HETRP* repeats, telomeric retrotransposons such as *HeT-A* are connected unidirectionally. **B**, Protein list with significant ChIP-Seq peaks on *HeT-A* as revealed by ChIP-Atlas (Oki S et al., 2018). Insulator proteins are highlighted with an inner rectangle. **C**, siRNA knockdown (KD) screen against insulator genes in OSC, followed by qRT-PCR of *HeT-A* transcript expression level. Values on y-axis are normalized by RP49 (n=3). Each dot indicates values obtained by different experiments. **D**, Volcano plot showing *HeT-A* expression level measured by qRT-PCR and corresponding significance upon KD using siRNA targeting 19 Mod(mdg4) variants and all variants. Three variants (variant N, V, AF) upregulating *HeT-A* upon their KD are labeled. Each dot represents KD of different variants (n=3). The x-axis is log2 ratio and the y-axis is log10 ratio. **E**, Western blotting (WB) showing protein levels of tubulin (loading control), total Mod(mdg4), Mod(mdg4)-N and HeT-A Gag upon KD of EGFP (control), Mod(mdg4)-AF+N+V, or Mod(mdg4) all variants. Molecular weight (MW) is indicated at the left of each image. **F**, WB showing protein levels of HeT-A Gag, Mod(mdg4)-N, total Mod(mdg4) and tubulin (loading control) in ovaries of +/+ wildtype (OregonR) or homozygous Mod(mdg4) RN1-2/RN1-2 mutant. Molecular weight (MW) is indicated at the left of each image. **G**, Bar plot showing results of crossbreeding experiments with indicated pairs of strains. RN1-2 and RN1-4 indicate lines bearing Mod(mdg4)-N variant-specific mutants (Fig. S2A). OregonR flies are used as wild type control. Left panel shows eggs per fly for 3 hours. Right panel shows hatching rate of eggs. N.A. indicates not applicable due to no eggs. **H**, Immunostaining images in adult ovaries from heterozygous (left) or homozygous (right) mutant flies of Mod(mdg4) RN1-2. These ovaries are stained with Orb (upper panels, red) or Vasa (lower panel, red) with DAPI (blue) staining. White bar indicates 50 µm. **I**, Immunostaining images of adult ovary in homozygous Mod(mdg4)-N mutant fly at low-power field (10×). The ovary is stained with Orb (red) and DAPI (blue). White bar indicates 200 µm.

*HeT-A* contains the Gag protein but not Pol, which includes the reverse transcriptase; therefore, *HeT-A* uses Pol from the other telomeric transposons upon its transposition (George et al., 2006; McGurk et al., 2021). Subtelomeres also have unique features. Subtelomere sequences of *D. melanogaster* are composed of subtelomeric *HETRP* (subtelomere heterochromatin repeat), also known as telomere-associated sequences, TAS (Karpen and Spradling, 1992) (Fig. 1A). Some of these repeat sequences have high similarity with the *Invader4* long terminal repeat (LTR) element (Bergman et al., 2006), suggesting a relationship between retrotransposons and telomere/subtelomere evolution. These unique features of telomere maintenance by retrotransposons in *D. melanogaster* look like a symbiosis between host and retrotransposon (Pardue and Debaryshe, 2002). However, de-repression of *HeT-A* retrotransposon in germ cells causes severe embryo abnormalities (Morgunova et al., 2015; Morgunova et al., 2021), and recent genomic analysis of the other *Drosophila* species has suggested that these retrotransposons have evolved very rapidly and replicated their copy number selfishly rather than symbiotically (Saint-Leandre et al., 2020; Saint-Leandre et al., 2019). Therefore, the host must regulate the activity of telomeric retrotransposons. In fact, multiple mechanisms to repress telomeric retrotransposons exist (Cacchione et al., 2020). For example, the PIWI-piRNA pathway is a small RNA-mediated RNA silencing mechanism (Iwasaki et al., 2015), known to repress telomeric retrotransposons in the fly ovary, and dysfunction of this pathway leads to intense *HeT-A* expression in germ cells (Savitsky *et al*., 2006; Senti et al., 2015; Shpiz et al., 2011; Teo et al., 2018). Heterochromatin machineries play an important role in regulating telomeric retrotransposons at some tissues as well. Heterochromatin protein 1 (HP1; also known as Su(var)205) mutation causes telomere fusions and increased expression of telomeric retrotransposons (Fanti et al., 1998; Perrini et al., 2004; Savitsky et al., 2002). Also, some transcription factors, such as *Woc, Hmr* or *Lhr*, and poly-A tail regulation machinery also contribute to *HeT-A* repression in ovaries (Font-Burgada et al., 2008; Morgunova *et al*., 2015; Raffa et al., 2005; Satyaki et al., 2014). However, even with these strong repressive machineries, developmental timing-specific and cell cycle-specific expression of telomeric retrotransposons are observed in wild-type flies (George and Pardue, 2003; Zhang et al., 2014), and the promoter of *HeT-A* is active when located in euchromatin (George and Pardue, 2003; Radion et al., 2017). These observations suggest that telomeric retrotransposons are regulated in a context-dependent manner with flexible regulatory machinery.

Chromatin insulators are sequences that possess enhancer blocking activities and act as epigenetic modification boundaries involved in the regulation of gene expression (Phillips-Cremins and Corces, 2013). These functions are mediated by the proteins bound to these insulator sequences. For example, the most characterized insulator protein so far is CTCF (CCCTC-binding factor). CTCF is necessary for enhancer-blocking and chromatin domain formation (Merkenschlager and Duncan, 2013; Rao et al., 2014), and most chromatin domain boundaries depend on CTCF and cohesin in vertebrates (Nora et al., 2017; Rao et al., 2017; Wutz et al., 2017). This is not the case in *Drosophila*, where a variety of insulator factors, including BEAF-32 and Su(Hw), among others, are responsible for insulator function and the *Drosophila* CTCF ortholog (dCTCF) has minor roles in chromatin conformation formation (Kaushal et al., 2021). Although the molecular basis of insulator function is not yet clear, earlier genetic studies proposed that the molecular mechanism responsible for the regulation of gene expression is controlled by loop formation between insulator elements (Cai and Shen, 2001; Melnikova et al., 2004; Muravyova et al., 2001).

Recently, Hi-C analysis has revealed that chromatin is divided into topologically associating domains (TADs), whose boundaries coincide with insulator proteins (Dixon et al., 2012; Lieberman-Aiden et al., 2009; Nora et al., 2012; Ramírez et al., 2018; Sexton et al., 2012; Wang et al., 2018). These observations suggest that insulators may be important for TAD formation. Although there is still much to clarify, recent studies have suggested that TADs are not necessarily required for enhancer-promoter interaction, but rather may play a role in assisting the enhancer function at least in *Drosophila* (Cavalheiro et al., 2021; Espinola et al., 2021; Ghavi-Helm et al., 2019; Ghavi-Helm et al., 2014; Ing-Simmons et al., 2021).

In this study, we show that specific splice variants of *Mod(mdg4)*, one of the insulator proteins in *Drosophila*, act as repressors of telomeric *HeT-A* retrotransposon by blocking subtelomeric enhancer activity in ovarian somatic cells (OSCs). By RNAi knockdown (KD) screening, we showed that Mod(mdg4) variants N, V, and AF (hereafter, Mod(mdg4)-N, V, AF respectively) are responsible for *HeT-A* repression.

Among these variants, Mod(mdg4)-N homozygous mutant flies display *HeT-A* de-repression in the ovary and female sterility phenotype. Further analysis revealed that each Mod(mdg4) variant has binding motif specificity and Mod(mdg4)-N is bound to both subtelomeric and telomeric loci in addition to genome-wide binding sites. Live-imaging analysis demonstrated enhancer blocking activity of subtelomeric *HETRP* repeats where Mod(mdg4)-N is bound, indicating that Mod(mdg4)-N represses *HeT-A* expression by blocking subtelomeric enhancers. Further investigation into how Mod(mdg4) blocks enhancer activities revealed the importance of its RNA polymerase II (Pol II) recruiting activity at subtelomeric repeats. For some of the Mod(mdg4)-N-binding sites other than telomeric or subtelomeric regions, we identified loss of chromatin boundaries upon depletion of Mod(mdg4)-N. Since these boundaries do not overlap with the sites where Pol II loss was observed upon Mod(mdg4)-N depletion, formation of chromatin boundaries by Mod(mdg4)-N is likely to be independent of Pol II regulation activities or *HeT-A* regulation. Our findings provide not only an unexpected subtelomere-telomere relationship mediated by insulator enhancer-blocking machinery, but also a conceptual framework of how insulator proteins orchestrate precise gene expression with different molecular mechanisms.

## Results

### Mod(mdg4) variants N, V, and AF repress *HeT-A* expression independently of heterochromatin formatio

*HeT-A* is repressed by the PIWI-piRNA pathway in the *Drosophila* ovary, where piRNAs against *HeT-A* are processed from Rhino-dependent transcription of piRNA precursor transcripts (Klattenhoff et al., 2009); therefore *HeT-A* is dramatically up-regulated in the ovary of *piwi* mutant flies. By contrast, piRNAs against *HeT-A* are not present in the cultured cell line driven from follicle cells of the *Drosophila* ovary, named OSC, where only Rhino-independent PIWI-piRNA regulation occurs (Saito et al., 2009). As a result, *HeT-A* up-regulation is not observed in *piwi* KD in OSC (Iwasaki et al., 2016) (Fig. S1A). KD of linker histone H1, which is indispensable for heterochromatin maintenance also does not cause up-regulation of *HeT-A* (Fig. S1A) (Iwasaki *et al*., 2016; Lu et al., 2009; Vujatovic et al., 2012), and levels of the heterochromatin histone mark H3K9me3 on *HeT-A* are relatively low compared to heterochromatic transposons including *mdg1* (Fig S1B). These results suggest the existence of a piRNA- and heterochromatin-independent *HeT-A* repression mechanism in OSC. To gain insight into the transcriptional regulation of retrotransposons at telomeres, we searched the ChIP-Atlas database for proteins associated to *HeT-A* coding regions according to published ChIP datasets (Oki et al., 2018). We found that several insulator proteins potentially associate to *HeT-A* coding regions (Fig. 1B). To test the possibility that *HeT-A* is regulated by these insulator proteins, we performed a KD screen. Mod(mdg4) was the only insulator protein to repress *HeT-A* expression in OSC (Fig.1C).

The *Mod(mdg4)* locus produces as many as 31 variants due to trans-splicing (Mongelard et al., 2002) (Fig. S1C). Among 31 variants, only variants H and T have had their function reported. Mod(mdg4)-T (also known as Mod(mdg4) 67.2) is a component of the gypsy insulator, which shows strong enhancer blocking activity (Cai and Levine, 1995; Gdula et al., 1996; Gerasimova et al., 1995). Lack of Mod(mdg4)-H (also known as Modifier of Mdg4 in Meiosis) leads to male infertility due to chromosome mis-segregation during meiosis (Sun et al., 2019; Thomas et al., 2005). Almost all variants were expressed in OSC with various expression levels (Fig. S1D). To determine which of the variants are responsible for the *HeT-A* repression, we performed a KD screen using variant-specific siRNAs. KD of variant N, V or AF [Mod(mdg4)-N, V, AF] resulted in up-regulation of *HeT-A* expression levels (Fig. 1D). Consistent with this observation, HeT-A Gag protein became detectable upon KD of these three variants (N+V+AF) to the level close to that observed upon KD of all variants (Fig. 1E, S1E-G). These results indicate that Mod(mdg4)-N, V, and AF repress *HeT-A* expression with piRNA- and heterochromatin-independent machinery.

### Mod(mdg4)-N mutant flies show elevated *HeT-A* expression and display female sterility phenotype

Mod(mdg4)-N KD resulted in the greatest degree of de-silencing of *HeT-A* expression among all tested variants in OSC (Fig. 1D). To investigate whether *HeT-A* up-regulation can be observed *in vivo* upon Mod(mdg4)-N loss and the physiological role of this specific variant, we generated two lines of Mod(mdg4)-N mutant flies (*RN1-2, RN1-4*) that harbor nucleotide deletions at the variant-specific exon using CRISPR-Cas9 (Kondo and Ueda, 2013) (Fig. S2A). Both deletions caused a frameshift in the coding sequence of the Mod(mdg4)-N-specific exon and resulted in loss of Mod(mdg4)-N protein in the homozygous mutant ovaries (Fig. 1F, S2B). In both homozygous mutant ovaries, we observed increased expression of both *HeT-A* transcript and HeT-A Gag proteins (Fig. 1F, S2B, C). This result indicates that Mod(mdg4)-N represses *HeT-A* expression *in vivo* as well as in OSC. Interestingly, flies homozygous for each mutant allele and trans-heterozygous flies were viable yet exhibited female sterility (Fig. 1G). *RN1-2* homozygous mutant flies lay eggs which, however, are morphologically abnormal and do not hatch (Fig. S2D), and *RN1-4* homozygous and trans-heterozygous mutant flies do not lay eggs at all. Although mutants of Mod(mdg4)-T or -H do not show such a female sterility phenotype, this phenotype resembles previously reported mutants that deplete all Mod(mdg4) variants (Büchner et al., 2000). Expression levels of variants other than Mod(mdg4)-N were not affected in this mutant (Fig. 1F, S2B), suggesting that Mod(mdg4)-N is responsible for the female sterility phenotype. Histologically, expression patterns of Orb, a marker for germ cells, and Vasa, a marker for germ cells and nurse cells, appeared normal at the early stages of oogenesis in Mod(mdg4)-N mutant flies (Fig. 1H). However, we observed residues of eggs in ovaries in Mod(mdg4)-N mutants (Fig. 1I), suggesting defective ovulation. De-repression of *HeT-A* in germ cells causes abnormal mitosis in early embryos and leads to lethality with maternal effect (Morgunova *et al*., 2015; Morgunova *et al*., 2021). However, these mutants lay eggs. In this context, Mod(mdg4)-N mutant flies show a more severe phenotype with impaired ovulation, which suggests that not only *HeT-A* up-regulation, but also other mechanisms contribute to the sterility phenotype. Altogether, these results indicate that Mod(mdg4)-N plays an important role in *HeT-A* repression and female fertility in the ovary.

### Mod(mdg4)-N specifically associates to telomeric and subtelomeric repeats

To uncover how Mod(mdg4)-N represses *HeT-A* expression, we performed ChIP-Seq analysis by generating stable cell lines expressing Ty1-tagged Mod(mdg4) variants (Fig. S3A).

Mod(mdg4)-T, which forms protein complexes with Su(Hw) and CP190 as a gypsy insulator, and binds to the same location as Su(Hw) (Nègre et al., 2010), was used as a control. To avoid artifacts caused by phantom or pseudo peaks (Jain et al., 2015), wildtype (WT) cells were also used to generate ChIP-Seq libraries. Of 31 Mod(mdg4) variants, 27 have FLYWCH domains in variant-specific regions (Dorn and Krauss, 2003). FLYWCH domains are the C2H2-type zinc finger domains conserved in eukaryotes, and necessary for DNA-binding or recruitment to chromatin (Beaster-Jones and Okkema, 2004; Melnikova et al., 2017), suggesting that genomic binding patterns of Mod(mdg4) variants may be different (Büchner *et al*., 2000). Consistent with this view, distribution of Mod(mdg4)-N and T on the genome showed variant-specific chromatin localization patterns and did not overlap (Fig. 2A, S3B, C). To characterize the sequence specificity of each Mod(mdg4) variant, we identified significantly enriched sequence motifs on their target sites. Peaks of each variant showed different sequence specificity as expected (Fig. 2B, S3D). The Mod(mdg4)-T motif was highly similar to the previously identified Su(Hw) motif (Fig. S3E), confirming that the result of Ty1 tag-based ChIP-Seq reflects the endogenous distribution of each Mod(mdg4) variant. The Mod(mdg4)-N motif resembled motifs of other transcription factors, NFAT (Nuclear Factor of Activated T cells) or Dl (Dorsal), but the similarity was not as high as that between Su(Hw) and Mod(mdg4)-T (Fig. S3E). Mod(mdg4)-V and AF also showed motif similarity with other transcription factors, but these transcription factors are not expressed at all in OSC (Fig. S3E). Furthermore, these Mod(mdg4) variants bound not only to the promoter regions where most transcription factors are enriched, but also to a variety of functional elements (Fig. 2C, S3F), suggesting that these Mod(mdg4) variants (other than Mod(mdg4)-T) recognize DNA sequences independently of other transcription factors.

**Fig. 2.**
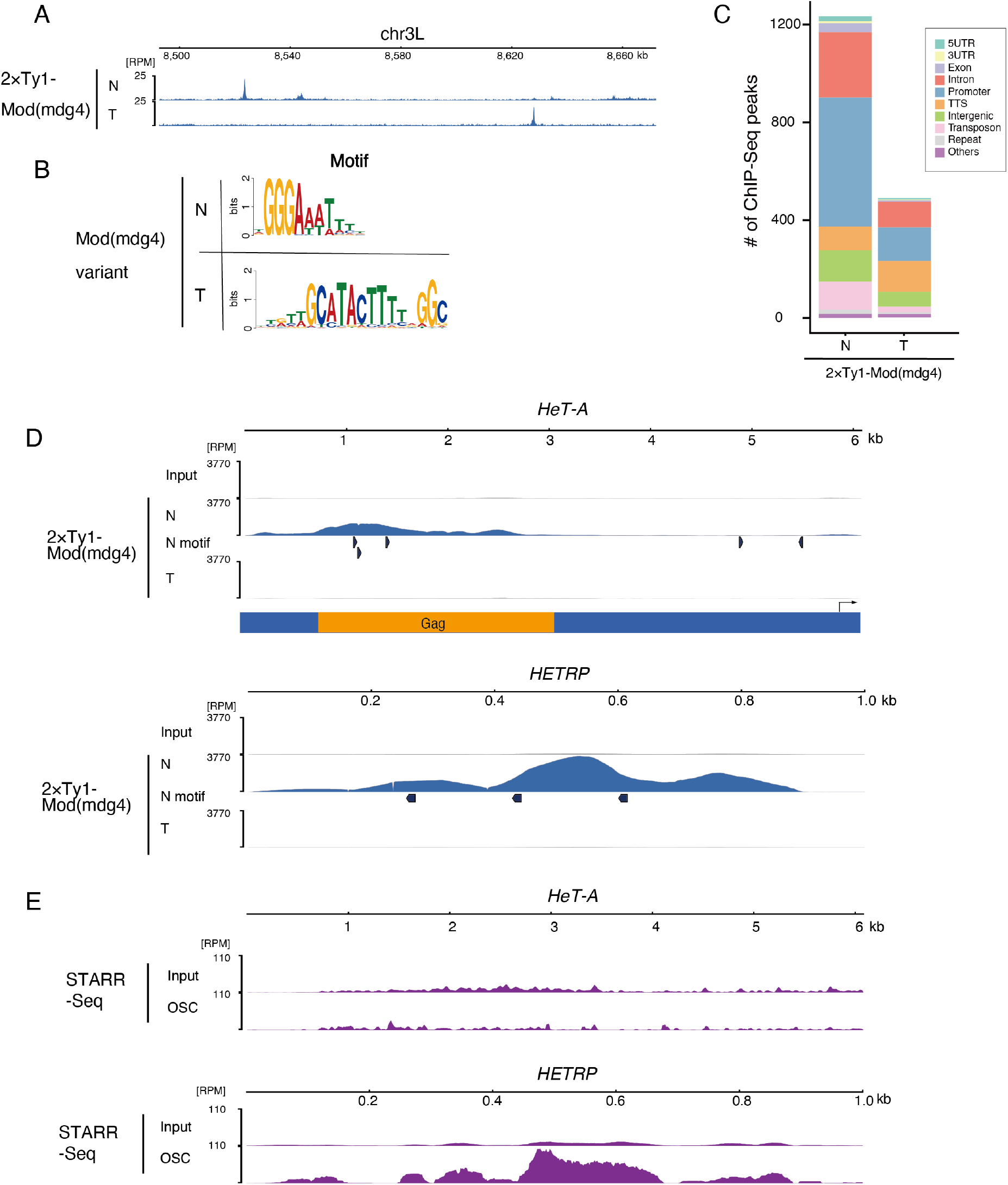
Mod(mdg4) recognizes different motifs by variant-dependent binding pattern. **A**, Example of distribution of each Mod(mdg4) variant. Coverage panel showing ChIP-Seq result of 2×Ty1-tagged Mod(mdg4) variants (N, T) at dm6 chr3L 8.50 Mb-8.66 Mb region. **B**, Logos showing clear enrichment around ChIP-Seq peaks of Mod(mdg4)-N and -T. These motifs are identified by de novo motif discovery analysis MEME (Multiple Em for Motif Elicitation). **C**, Functional annotation of peaks of Mod(mdg4)-N and -T, identified by HOMER. The y-axis shows number of peaks of each Mod(mdg4) variant. We used only highly confident (FDR < 1e-15) ChIP-Seq peaks for analysis to avoid ambiguous peaks. **D**, Distribution of Mod(mdg4)-N and -T on consensus sequence of HeT-A (upper panel) and HETRP (lower panel) from repbase. Coverage of each ChIP-Seq experiment is shown with blue, and motifs of each Mod(mdg4)-N are indicated as black arrows below the coverage panels. The y-axis is normalized with RPM. Below coverage panel of HeT-A, the reading frame of HeT-A Gag protein is shown in orange, and the black arrow indicates the HeT-A promoter. **E**, Coverage panel showing STARR-Seq signal (purple) for OSC and its input on consensus sequence of HeT-A (upper panel) and HETRP (lower panel) from repbase. The y-axis is normalized with RPM. This data is reanalyzed from STARR-Seq results of Arnold CD et al., 2013.

We next investigated how Mod(mdg4)-N associates to telomere/subtelomere sequences. As expected from the KD screen (Fig. 1C), mapping to the consensus sequence of *HeT-A* revealed that Mod(mdg4)-N is bound to *HeT-A* whereas Mod(mdg4)-T is not (Fig. 2D; upper panel). Additionally, Mod(mdg4)-N bound to *HETRP* repeats strongly (Fig. 2D; lower panel), and the motif enriched on Mod(mdg4)-N peaks was observed on *HeT-A* or *HETRP* sequences (Fig. 2D). Taken together, accumulation of Mod(mdg4)-N on telomeric *HeT-A* and subtelomeric *HETRP* repeats suggests that the variant directly regulates expression of *HeT-A* at telomere and subtelomere loci rather than by regulating other genes.

Previous reports suggested enhancer-blocking activity of Mod(mdg4)-T as a component of the *gypsy* insulator (Gerasimova *et al*., 1995; Kyrchanova et al., 2013). Taking this into account, the strong binding of Mod(mdg4)-N on the telomeric and subtelomeric repeat sequences suggests that Mod(mdg4)-N limits access of enhancer to *HeT-A* promoter and induces silencing of *HeT-A*. To test this, we re-analyzed STARR-seq [self-transcribing active regulatory region (STARR) sequencing] data, which assess enhancer activity by parallel reporter plasmid in OSC (Arnold et al., 2013), and searched for enhancer sequences on *HeT-A* or *HETRP*.

Interestingly, *HETRP* had strong enhancer activity in OSC, whereas enhancer activity was not observed on *HeT-A* (Fig. 2E). It is plausible that *HETRP* repeats possess enhancer activity because *HETRP* repeats are derived from LTR sequences of the *Invader4* retrotransposon (Asif-Laidin et al., 2017; Bergman *et al*., 2006), and because LTR sequences generally have both enhancer and promoter activity (Thompson et al., 2016). With these observations on Mod(mdg4)-N-binding features and cis-regulatory element landscape, we hypothesized that Mod(mdg4)-N blocks subtelomeric enhancers to suppress *HeT-A* expression.

### Subtelomeric *HETRP* repeats possess enhancer-blocking activity

To test this hypothesis, we performed live-imaging analysis of enhancer-blocking activities of Mod(mdg4)-N-binding sequences on telomere/subtelomere in the living embryo, where Mod(mdg4) expresses at a high level (Büchner *et al*., 2000). In this imaging system, *sna shadow* enhancer causes a transcriptional burst from the *Drosophila* synthetic core promoter (DSCP), which transcribes 24xMS2-yellow synthetic gene. MCP (MS2 coat protein) is a single-stranded RNA phage capsid protein that binds to the MS2 19-nucleotide RNA stem loop with high affinity (Bertrand et al., 1998), and MCP-GFP fusion protein recognizes MS2 repeats on this synthetic gene. With this imaging system, transcriptional dynamics and enhancer function can be visualized precisely (Fukaya et al., 2016; Garcia et al., 2013; Lucas et al., 2013) (Fig. 3A). For the enhancer blocking assay, we tested two Mod(mdg4)-N-binding *HETRP* and *HeT-A* sequences (Fig. S4A). We compared these sequences with gypsy insulator sequence, which strongly inhibits enhancer-promoter interaction via gypsy insulator proteins (Bender et al., 1983; Fukaya *et al*., 2016; Ghosh et al., 2001). Although a single insertion of *HeT-A* sequence between enhancer and promoter did not affect transcriptional bursting frequency nor amplitude of bursting, a single insertion of *HETRP* decreased transcriptional bursting frequency (Fig. 3B-E, Supplementary Video 1), indicating that the *HETRP* sequence is a *bona fide* insulator. Amplitude per burst did not change (Fig. 3F,G). Considering that Mod(mdg4)-N mutants display 5–10-fold higher *HeT-A* expression in ovary (Fig. S2C), this enhancer-blocking activity of a single *HETRP* is weaker than expected. Both *HETRP* and *HeT-A* exist as tandem repeats in the *D. melanogaster* genome (Karpen and Spradling, 1992; Pardue and Debaryshe, 2002). Therefore, we reasoned that mimicking *in vivo* tandem repeats of *HETRP* and *HeT-A* is important to assess enhancer-blocking activity of these sequences. We further inserted *HETRP* sequences at both sides of the *sna shadow* enhancer.

**Fig. 3.**
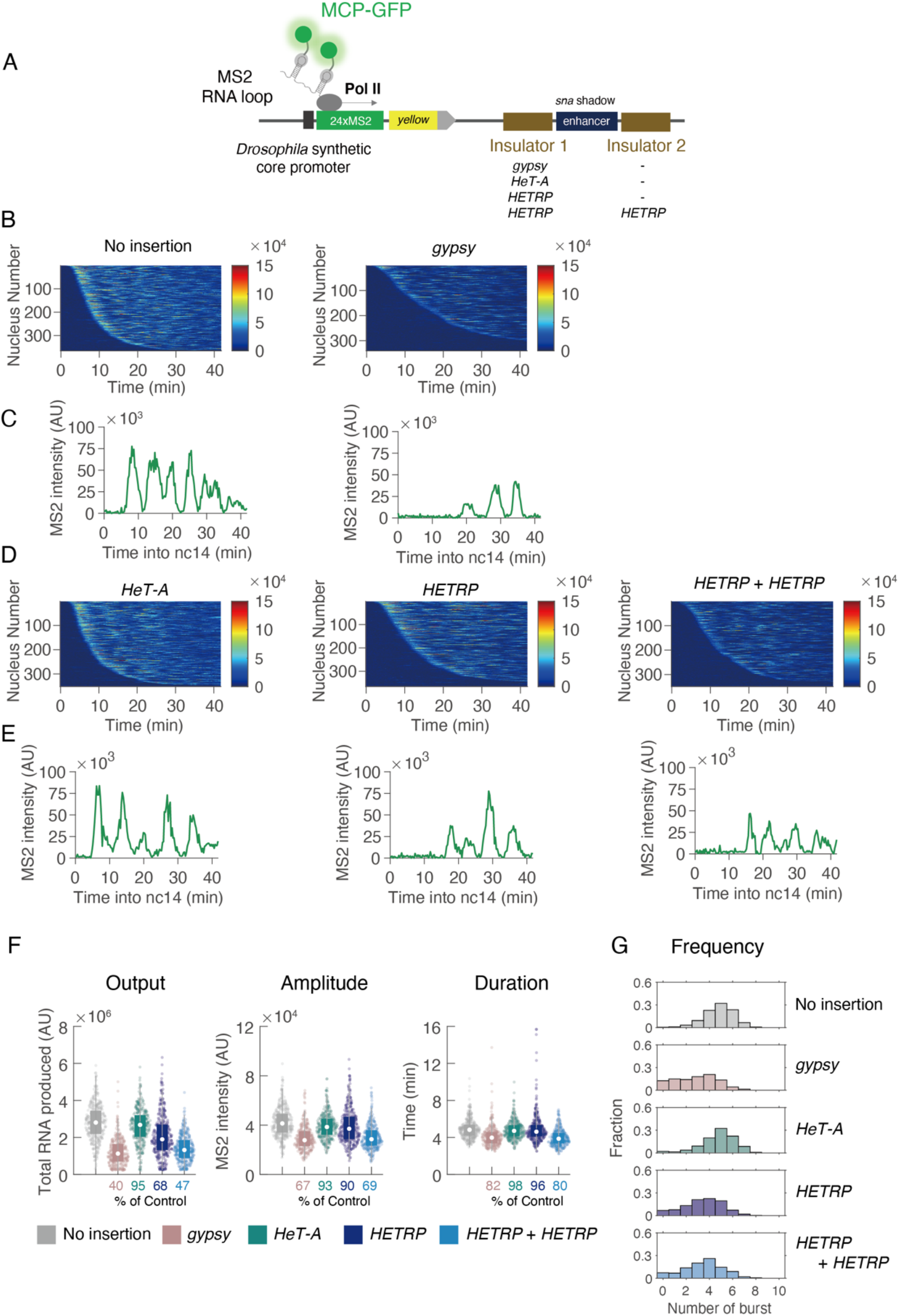
Subtelomeric *HETRP* repeats possess enhancer-blocking activity. **A**, Schematic representation of the yellow reporter gene containing the 155-bp *Drosophila* synthetic core promoter (DSCP), the 1.5-kb sna shadow enhancer, and 24× MS2 RNA stem loops within the 5′ UTR. Insulator candidate sequences are inserted at both sides of *sna shadow* enhancer. Expression of this reporter gene is visualized by MCP-GFP proteins binding to MS2 RNA stem loops. **B**, MS2 trajectories for all analyzed nuclei. Each row represents the MS2 trajectory for a single nucleus. A total of 365 and 340 ventral-most nuclei, respectively, were analyzed from three independent embryos for the reporter genes with no insertion of insulators (left), or *gypsy* insulator (right). Nuclei were ordered by their onset of transcription in nc14. AU; arbitrary unit. **C**, A representative trajectory of transcriptional activity of the MS2 reporter gene with control (left), or single insertion of *gypsy* insulator (right). **D**, MS2 trajectories for all analyzed nuclei. Each row represents the MS2 trajectory for a single nucleus. The ventral-most nuclei were analyzed from three independent embryos for the reporter genes with single insertion of *HeT-A* sequence (left), single insertion of *HETRP* sequence (middle) and double insertion of *HETRP* sequence (right). A total of 352, 365 and 349 ventral-most nuclei, respectively, were ordered by their onset of transcription in nc14. AU; arbitrary unit. **E**, A representative trajectory of transcriptional activity of the MS2 reporter gene with single insertion of *HeT-A* sequence (left), single insertion of *HETRP* sequence (middle) and double insertion of *HETRP* sequence (right). **F**, Boxplots showing the distribution of total output (left), burst amplitude (middle) and burst duration (right). The box indicates the lower (25%) and upper (75%) quantile and the open circle indicates the median. Whiskers extend to the most extreme, non-outlier data points. A total of 365, 340, 352, 365 and 349 ventral-most nuclei, respectively, were analyzed from three independent embryos for the reporter genes with no insertion of insulators, single insertion of *gypsy* insulator, *HeT-A* sequence, or *HETRP* sequence and double insertion of *HETRP* sequence, from left to right. Median values relative to the control reporter are shown at the bottom. AU; arbitrary unit. **G**, Histograms showing the distribution of burst frequency during nc14 stage. A total of 365, 340, 352, 365 and 349 ventral-most nuclei, respectively, were analyzed from three independent embryos for the reporter genes with no insertion of insulators, single insertion of *gypsy* insulator, *HeT-A* sequence, or *HETRP* sequence and double insertion of *HETRP* sequence, from top to bottom.

Two copies of *HETRP* sequence resulted in strong blocking of the enhancer activities compared with a single insertion (Fig. 3D-E). With two copies of *HETRP*, not only frequency of transcriptional bursting but also amplitude and duration per burst decreased (Fig. 3F-G). This result suggests that enhancer-blocking is specific to the *HETRP* sequence, and multiple repeat alignment in the genome additively facilitates enhancer-blocking activity of *HETRP* sequences. By contrast, two copies of *HeT-A* changed neither total output, amplitude nor frequency (Fig. S4B-E, Supplementary Video 2). These results indicate that Mod(mdg4)-N-binding *HETRP* sequences are bona fide insulators, and multiple repeat alignment of *HETRP* is important to block the enhancer activity within *HETRP* sequences.

### RNA polymerase II is recruited to Mod(mdg4)-N binding sites

Next, we dissected molecular details about how Mod(mdg4)-N works at its binding sites. Previous studies have demonstrated a link between RNA Pol II and insulation: enhancer effect on a promoter is blocked when promoters are inserted between them (Chopra et al., 2009; Kaushal *et al*., 2021) and promoters with strong Pol II accumulation interact with insulator proteins (Liang et al., 2014). Consistent with these observations, approximately half of the Mod(mdg4) peaks bound to promoters (Fig. 2C). It is plausible that Mod(mdg4)-N acts as an insulator by regulating Pol II accumulation. Promoters with high promoter-proximal Pol II pausing block enhancer-promoter communication (Chopra *et al*., 2009). To analyze the relationship between Mod(mdg4)-N and promoter-proximal pausing, we first reanalyzed public data for OSC Pol II ChIP (Sienski et al., 2012) using the pausing index (Fig. S5A). The pausing index is the ratio of the amount of Pol II at the region 250 bp either side of the TSS to the amount of Pol II from TSS+500 bp to TTS-500 bp (Gilchrist et al., 2010). The higher this index, the higher the pausing observed at the promoter. The promoters with Mod(mdg4)-N had a significantly higher pausing index when compared to all promoters (Fig. S5B; Kolmogorov-Smirnov test p=2.22×10^−11^), showing that Mod(mdg4)-N tends to bind to highly pausing promoters. We next examined how the distribution of Pol II changes upon Mod(mdg4)-N KD. Mod(mdg4)-N bound to only selected promoters; therefore, global changes in Pol II accumulation were not observed upon Mod(mdg4)-N KD. However, some Pol II peaks were significantly lost (log_2_(fold change) < -0.1) upon Mod(mdg4)-N KD (Fig. 4A). By contrast, almost no Pol II peaks became more pronounced upon Mod(mdg4)-N KD. These ‘Pol II loss genes’ were also observed in KD of all Mod(mdg4) variants and highly overlapped with Mod(mdg4)-N peaks (Fig. 4B, C).

**Fig. 4.**
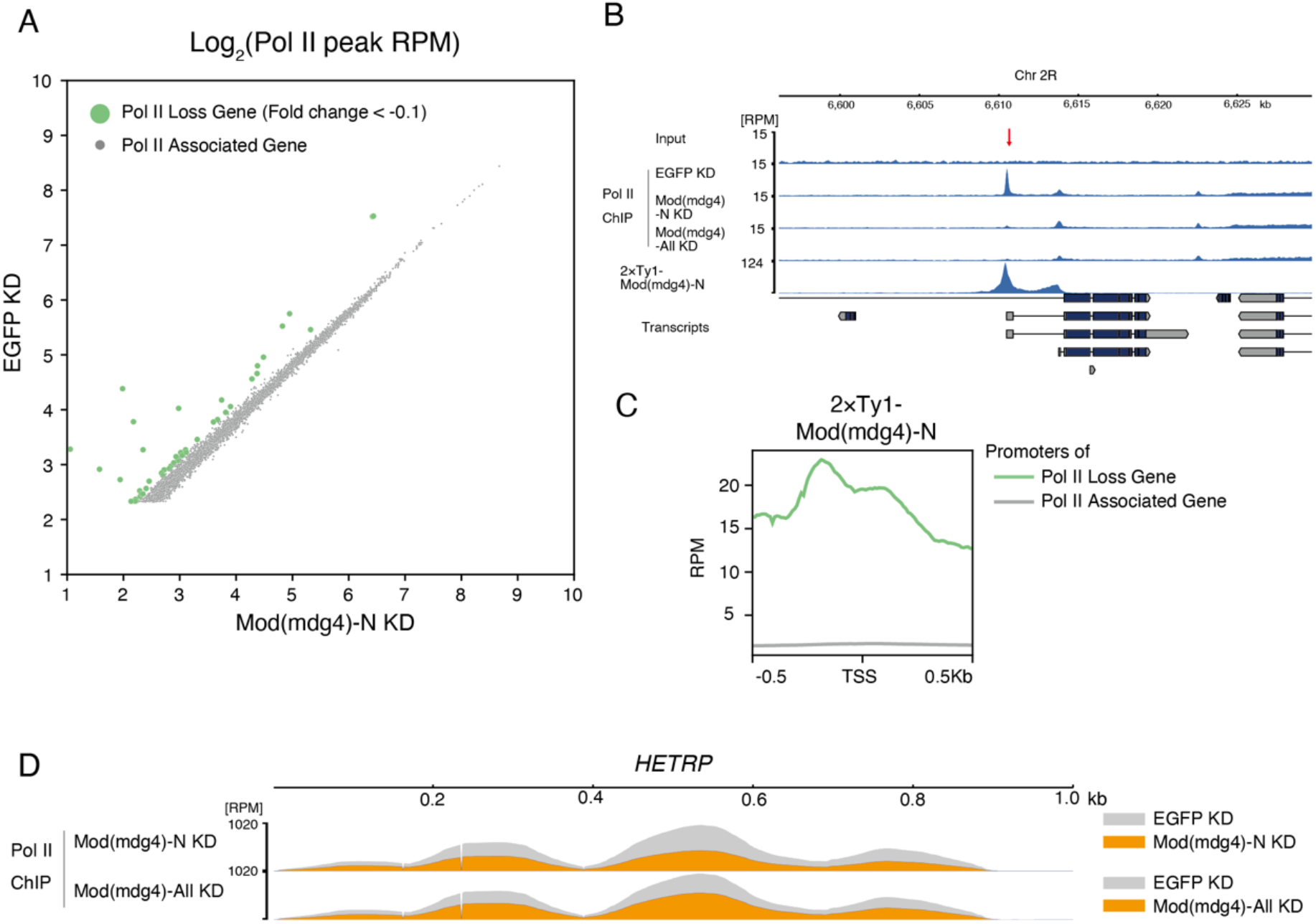
RNA polymerase II is recruited to Mod(mdg4)-N binding sites. **A**, Scatter plot of peak maximum coverage between EGFP KD and Mod(mdg4)-N KD. Both x-axis and y-axis are shown with log2 ratio. Peaks whose maximum values are under 5.0 at RPM are removed from analysis. Pol II Loss genes (green) are defined as log2(Fold Changes) < -0.1 and other genes (Pol II Associated genes) whose Pol II level does not change upon Mod(mdg4)-N KD are shown with gray dots. **B**, Coverage tracks showing localization of Mod(mdg4)-N and pol II profile upon EGFP KD, Mod(mdg4)-N KD and all variant KD at Pol II loss gene locus (Chr2R:6.60Mb-6.63Mb). Lower panel indicates annotation of transcripts from Flybase. A red arrow indicates the pol II peak that disappeared upon Mod(mdg4)-N or all variants KD. **C**, Average profile showing enrichment of Mod(mdg4)-N ChIP-Seq intensity around Pol II peaks that disappear upon Mod(mdg4)-N KD (Pol II Loss genes; green) or the other genes (Pol II Associated gene, gray). **D**, Coverage tracks showing Pol II enrichment on consensus sequence of subtelomeric *HETRP* repeats upon EGFP KD, Mod(mdg4)-N KD, or Mod(mdg4) all variants KD. ChIP signal depths on EGFP KD (gray), Mod(mdg4)-N KD (orange, upper panel) and Mod(mdg4) all variant KD (orange, lower panel) are shown.

Surprisingly, the *HETRP* repeats at subtelomeres were also associated with Pol II, and KD of Mod(mdg4)-N or all variants of Mod(mdg4) resulted in loss of Pol II from this region (Fig. 4D). Sequence analysis revealed that core promoter sequences, Initiator (Inr) (Purnell et al., 1994) and Downstream Promoter Element (DPE) (Burke and Kadonaga, 1996), for Pol II association exist in a *HETRP* repeat with the criteria described in previous research (Shao et al., 2019) (Fig. S6A). This observation implies that Pol II regulated by Mod(mdg4)-N is important for enhancer blocking activity. To test this, we investigated whether loss of *HETRP* core promoter sequence attenuates its enhancer blocking activity in OSC. We designed a reporter where the *Tj* enhancer acts on DSCP, which expresses EGFP:P2A:BlastR, and assessed *Tj* enhancer blocking activity of *HETRP* mutant sequences by EGFP intensity (Fig. S6B).

This reporter vector was integrated into chromatin with the PiggyBac system (Fig. S6C), as chromatin integration is known to improve reproducibility of enhancer reporter assays (Inoue et al., 2017). To analyze the impact of Pol II association on enhancer blocking activity, we compared two *HETRP* sequence mutants for enhancer reporter assay: the first mutant was a complete deletion of core promoter sequences, Inr and DPEs, from a *HETRP* sequence; the second mutant was a replacement of the Inr sequence with a random sequence (Fig. S6A). With this reporter assay, we observed enhancer blocking activity of two copies of *HETRP* repeats (Fig. S6D), consistent with the live imaging assay (Fig. 3). Both deletion and replacement of *HETRP* core promoter sequences weakened the enhancer blocking activity, though these mutations did not cause complete loss of enhancer blocking activities. Thus, Pol II recruited on *HETRP* repeats assists enhancer blocking by Mod(mdg4)-N.

To further investigate Pol II accumulation observed in Mod(mdg4)-N-associated regions, we tested whether this is due to promoter-proximal pausing. Promoter-proximal pausing controls expression of downstream genes and requires two steps: recruitment of preinitiation complex and Pol II, which positively regulate transcription, and recruitment of DSIF (DRB Sensitivity Inducing Factor) and NELF (negative elongation factor), which negatively regulate transcription (Core and Adelman, 2019). However, depletion of proteins necessary for promoter-proximal pausing did not affect *HeT-A* expression levels (Fig. S6E), suggesting that Mod(mdg4)-N functions at the recruitment step of Pol II regulation but not at the step of promoter-proximal pausing for *HeT-A* regulation. Overall, these results suggest that recruitment of Pol II by Mod(mdg4)-N assists *HeT-A* repression by blocking subtelomeric enhancer, and enhancer blocking activity by Mod(mdg4)-N is the major mechanism regulating *HeT-A* expression.

### Mod(mdg4)-N affects formation of chromatin domain boundaries independently of Pol II regulation

Mod(mdg4)-N showed genome-wide distribution (Fig. 2). Most Mod(mdg4)-N associated regions were not tandemly repeated as in telomeric regions, and not all Mod(mdg4)-N-binding promoters showed Pol II loss upon Mod(mdg4)-N KD (Fig. 4A). These observations suggest that Mod(mdg4)-N associated to non-telomeric regions has functions other than enhancer blocking. To investigate this, we focused on higher-order chromatin conformation because insulator proteins including Mod(mdg4) frequently localize at TAD boundaries and affect higher-order chromatin conformation in *Drosophila*, although there are some inconsistences among studies as to which insulator proteins accumulate on TAD boundaries (Cubeñas-Potts et al., 2017; Ramírez *et al*., 2018; Sexton *et al*., 2012; Wang *et al*., 2018).

To investigate how Mod(mdg4)-N affects higher-order chromatin conformation with finer resolution, we performed Micro-C XL, a variation on Hi-C, using OSC (Hsieh et al., 2020; Hsieh et al., 2016; Krietenstein et al., 2020). We generated a 200-bp resolution contact map with detailed contact information especially at short range (Fig. 5A, S7A) (Lieberman-Aiden *et al*., 2009; Naumova et al., 2013; Wang *et al*., 2018), which enabled us to analyze small chromatin domains (median size of 7 kb at 200 bp bin size) (Fig. S7B).

**Fig. 5.**
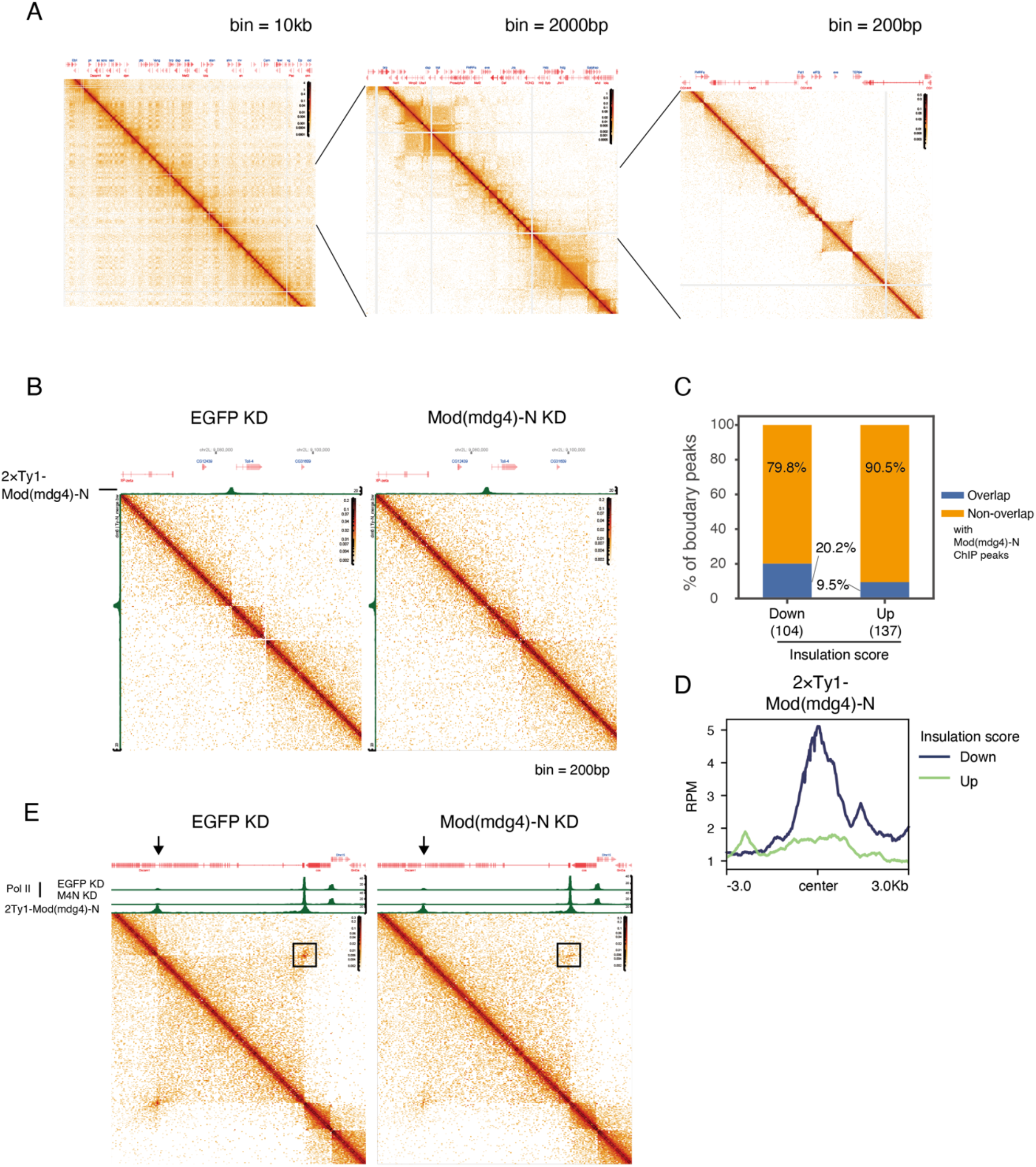
Mod(mdg4)-N forms topological domain boundaries on its binding sites. **A**, Micro-C XL contact map near dm6 chr2R:9.5-10.5Mb at different resolution (10 kbp, 2000 bp and 200 bp). Gene annotation from Flybase is shown at the top of each panel. **B**, Heat map showing example of region (near Toll-4 gene) where loss of Mod(mdg4)-N causes insulation loss. Each panel shows contact heat map upon EGFP (control) KD (left) and Mod(mdg4)-N KD (right). ChIP-Seq signal of Mod(mdg4)-N is indicated at the top of each panel (green). **C**, Percentage of sites overlapping with Mod(mdg4)-N ChIP peaks where insulation score called at 200 bp resolution decreases (104 sites, left) and increases (137 sites, right) upon Mod(mdg4)-N KD. **D**, Average profile showing Mod(mdg4)-N ChIP-Seq enrichment around the sites where insulation score decreases (blue) and increases (green) upon Mod(mdg4)-N KD over ±3.0 kb region from boundaries. **E**, Contact map of EGFP KD (left panel) and Mod(mdg4)-N KD (right panel) showing region where loss of Mod(mdg4)-N cause genomic looping loss. Black rectangles indicate looping between Dscam1 promoter and intron region. Coverage tracks of Mod(mdg4)-N and Pol II distribution upon EGFP (control) and Mod(mdg4)-N KD are shown at top of contact map.

Using this contact map, we analyzed co-localization between Mod(mdg4) peaks and TAD boundaries. Half of Mod(mdg4)-N peaks (49.1%) co-localized with domain boundary detected at 200 bp bin size (Fig. S7C). This percentage was slightly lower than that for CP190 (Centrosomal protein 190kD) (66.1%), which functions as a central component of *Drosophila* chromatin conformation (Ramírez *et al*., 2018; Wang *et al*., 2018), and Mod(mdg4)-T (72.4%). However, the overlapping rate between Mod(mdg4)-N and chromatin boundaries was much higher than random sequence (49.1% vs 7.7%), suggesting strong chromatin boundary preference of Mod(mdg4)-N localization.

These co-localization patterns led us to ask whether disruption of Mod(mdg4)-N-binding causes chromatin conformation change. We performed Micro-C XL in Mod(mdg4)-N KD and observed chromatin conformation change. For some of the Mod(mdg4)-N associated regions, overlapping boundary was lost (Fig. 5B), suggesting the loss of insulation. Genome-wide, we observed 104 sites with insulation loss and 137 sites with insulation gain upon Mod(mdg4)-N KD (Fig. 5C). Importantly, 20.2% of sites with insulation loss overlapped with Mod(mdg4)-N-binding sites and clearly showed center enrichment of Mod(mdg4)-N, whereas only 9.5% of sites with insulation gain overlapped with Mod(mdg4)-N-binding sites and did not show Mod(mdg4)-N center enrichment (Fig. 5C,D). These results suggest that Mod(mdg4)-N functions mainly at regions where insulation is lost upon Mod(mdg4)-N KD and thus, Mod(mdg4)-N has a role in establishment of domain boundaries on the genome at its binding site. At these sites with insulation loss, we observed no significant change of Pol II distribution (Fig. S7D). This result suggests that Mod(mdg4)-N regulates formation of chromatin domains on its binding sites, and this is independent of the regulation mechanism observed at the telomere and subtelomere.

Although Pol II recruitment functions of Mod(mdg4)-N are not linked to chromatin domain formation, Mod(mdg4) is one of the proteins enriched at chromatin loop anchors among insulator proteins (Cubeñas-Potts *et al*., 2017). Interestingly, some sites showing loss of looping signature overlapped with sites that lost Pol II upon Mod(mdg4)-N KD. It is known that a genomic loop is shown as a dot in the contact map (Hsieh *et al*., 2020; Krietenstein *et al*., 2020), and we observed dot-like structures between two Mod(mdg4)-N bound sites at at least two loci (*Dscam1* promoter-intron loop and *comm* promoter – *comm2* promoter) (Fig. 5E, S7E; black rectangles). In Mod(mdg4)-N KD, dot signatures weakened at these loci and Pol II loss was observed, indicating that formation of these loops depends on Mod(mdg4)-N Pol II recruiting activity. Whether similar looping by Mod(mdg4)-N occurs at subtelomeric *HETRP* regions remains to be established, because the highly repetitive nature of the telomere/subtelomere regions makes it difficult to analyze Hi-C data. However, we observed Pol II loss at subtelomeric *HETRP* regions upon Mod(mdg4)-N KD (Fig. 4D), suggesting that loop formation by Mod(mdg4)-N might play an important role in enhancer-blocking at subtelomeric *HETRP* regions, rather than TAD boundary formation. Overall, these results suggest that Mod(mdg4)-N plays multiple roles in chromatin regulation, possibly in a context-dependent manner. Mod(mdg4)-N forms TAD boundaries independently of its Pol II regulating activity, possibly at single associated sites.

Meanwhile, changes in the loop formation associated with Pol II accumulation might play an important role in subtelomeric enhancer-blocking, because enhancer-blocking activity of subtelomeric *HETRP* repeats is facilitated when the repeats are tandemly aligned (Fig. 3).

## Discussion

In this study, we showed that a specific Mod(mdg4) variant represses *HeT-A* retrotransposon by blocking of enhancer on subtelomeric repeats along with Pol II recruitment. Moreover, we observed changes in chromatin conformation upon Mod(mdg4)-N depletion in a mechanism independent of its Pol II regulating activity. These effects on chromatin may function to orchestrate proper gene expression.

### Enhancer and insulator function of *Drosophila* subtelomere

We propose a model in which the enhancer function of the *HETRP* region is blocked by Mod(mdg4)-N, which in turn represses the expression of *HeT-A* at telomeres (Fig. 6A). We observed drastic Pol II loss on *HETRP* upon Mod(mdg4)-N depletion (Fig. 4D), and deletion or mutation of core promoter sequences on *HETRP* attenuated but did not completely abolish its insulator activity (Fig. S6D). With this observation, we propose that two mechanisms are involved in the enhancer blocking function of Mod(mdg4)-N. The first is Pol II recruitment-dependent insulator activity as we have shown (Fig. S6). Possibly, these transcription machineries modulate insulator function by open chromatin formation to recruit additional factors to subtelomeric regions (Gilchrist *et al*., 2010; Shopland et al., 1995), or by loop formation between promoters on *HETRP* repeats (Fig. 5E). The second is the biochemical nature of insulator proteins. Mod(mdg4) has a BTB/POZ domain at its N-terminal common region, which can form multimers with both itself and a BTB/POZ domain of Trl (Trithorax-like, a.k.a. GAGA Factor) (Bonchuk et al., 2011; Melnikova *et al*., 2004). We suggest that this homophilic nature of a Mod(mdg4) BTB/POZ domain contributes to enhancer-blocking by dynamic loop formation (Fukaya *et al*., 2016; Kyrchanova *et al*., 2013).

**Fig. 6.**
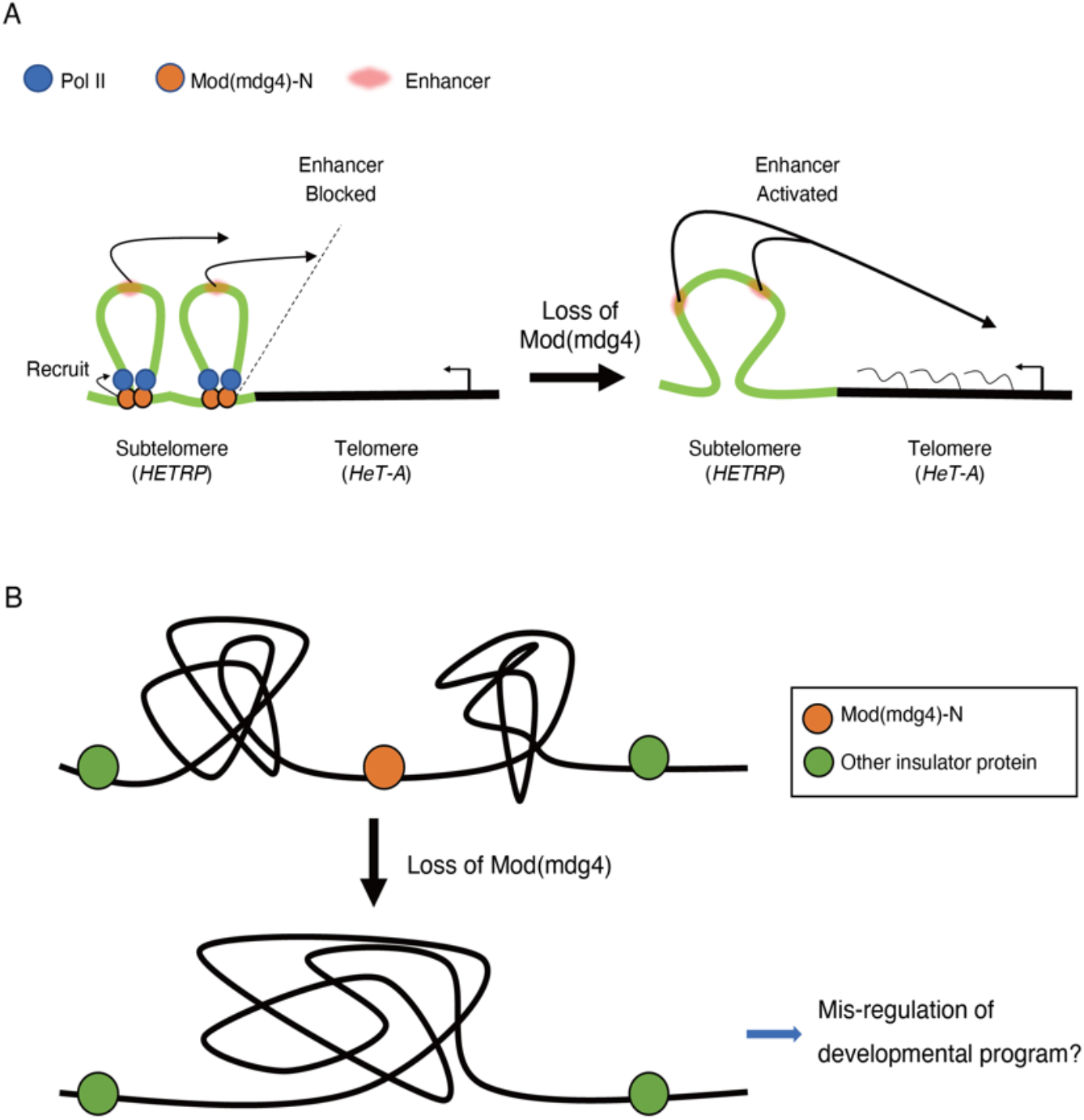
Summary of this research. **A**, Mod(mdg4)-N (orange) recruits pol II (blue) at subtelomeric region, which assists blocking of subtelomeric enhancer (red). Therefore, *HeT-A* is silenced in the normal state. Upon loss of Mod(mdg4), enhancer-blocking activity is lost, leading to *HeT-A* expression. **B**, Function of Mod(mdg4)-N in forming chromatin boundary. Loss of Mod(mdg4)-N causes fusion of these domains, which leads to mis-regulation in development.

By contrast, we could not show enhancer-blocking activity of the *HeT-A* sequence, although Mod(mdg4)-N binds to this sequence (Fig. 2D, 3, S4). There are two non-exclusive scenarios to explain this observation. The first scenario is the presence of unknown trans-acting factors. Enhancer-blocking activity is highly context-dependent. For example, well-characterized *gypsy* insulator is composed of Su(Hw), CP190 and Mod(mdg4)-T. However, not all of three main gypsy insulator component binding sites show enhancer blocking activity (Nègre et al., 2011; Schwartz et al., 2012). Some other trans-acting factors that do not exist on *HeT-A*, such as promoter machinery as we showed (Fig. S6D), might be necessary for enhancer-blocking activity. The other scenario is simply that affinity of Mod(mdg4)-N-binding to telomeric *HeT-A* is weaker than that to *HETRP* repeats (Fig. 2D). Although we could not show a direct role of Mod(mdg4)-N on *HeT-A* in this study, we speculate that Mod(mdg4)-N might also play an important role such as TAD boundary formation or epigenetic barrier activity in a specific context (Phillips-Cremins and Corces, 2013). The repetitive nature of these sequences makes it difficult to analyze affinity or 3D structural impact *in vivo*. To analyze these unique sequences, further technical improvements that can be used for repeated sequences will be needed.

Overall, our results indicate that Mod(mdg4)-N is necessary for *HeT-A* repression by blocking subtelomeric enhancer. Interestingly, enhancer-like function of subtelomere is also reported in human, and transcription of TERRA (telomeric repeat-containing RNA) is promoted by insulator protein CTCF despite telomeric sequences in human and fly being completely different (Deng et al., 2012; Tutton et al., 2016). This observation shows an evolutionarily conserved principle of enhancer-like functions at subtelomeres being modulated by insulator proteins. It is still an open question what drives this subtelomeric convergent evolution.

### Complexity and specificity of Mod(mdg4) variants

Mod(mdg4) has 31 variants that are generated a by trans-splicing mechanism (Mongelard *et al*., 2002). We showed that binding sites of each Mod(mdg4) variant rarely overlap, and each variant is enriched at different motifs (Fig. 2, S3).

Functionally, we observed that the Mod(mdg4)-N mutant displays a female sterility phenotype. This was observed in the mutant of all Mod(mdg4) variants, but not reported as the phenotype of the other specific Mod(mdg4) variants, underlining the uniqueness of Mod(mdg4)-N function. These results indicate that different chromatin localization patterns are responsible for different physiological roles *in vivo*. Mod(mdg4) has a BTB/POZ domain at its N-terminal common region, whereas most of the C-terminal variant-specific regions contain FLYWCH zinc finger domains (Dorn and Krauss, 2003). These domain structures suggest that the BTB/POZ domain is important for formation of chromatin conformation via its self-interacting nature (Bonchuk *et al*., 2011), and FLYWCH domains define the sequence specificity of chromatin binding. Although the FLYWCH domain of Mod(mdg4)-T is responsible for its interaction with Su(Hw) (Ghosh *et al*., 2001), the FLYWCH domain in *Caenorhabditis elegans* PEB-1 directly binds to DNA (Beaster-Jones and Okkema, 2004). The motifs enriched at Mod(mdg4)-N-binding sites did not have high similarity with any of the known transcription factor motifs (Fig. 2B), and Mod(mdg4) variants bind on variously annotated sites, not only on promoters. With these observations and previous studies, it is likely that Mod(mdg4)-N directly binds to DNA via its FLYWCH domain.

### Insulator functions and multiple mechanisms of regulation by Mod(mdg4)

We showed that Mod(mdg4)-N functions as an insulator, but its role is different from Mod(mdg4)-T, a component of the gypsy insulator. This means that Mod(mdg4) acts in a variant-specific manner, and the list of insulator proteins (e.g., CP190, BEAF-32, Su(Hw), dCTCF) in *Drosophila* may be expanded. In contrast to *Drosophila*, mammals have only one main insulator protein, CTCF (Nora *et al*., 2017; Phillips and Corces, 2009). Loss of some insulator proteins results in a tissue-specific phenotype rather than embryonic lethality in *Drosophila* (Gambetta and Furlong, 2018; Kaushal *et al*., 2021), and we also showed that loss of Mod(mdg4)-N caused a sterile phenotype but did not affect viability (Fig. 1). This is also in contrast to mammals, where loss of CTCF causes severe embryonic defects (Fedoriw et al., 2004). So, how do insulator proteins orchestrate proper gene expression in *Drosophila*? In this study, we showed bi-functionality of Mod(mdg4)-N; Pol II recruitment activity (Fig. 4) which assists subtelomeric enhancer blocking, and formation of chromatin boundaries (Fig. 5) as an independent mechanism. This means that TAD formation is only one of the functions of insulator proteins.

Although the function of chromatin conformation or TADs is under debate, TADs are likely to have a supporting role to ensure proper gene expression rather than being necessary factors for enhancer-promoter communication (Ghavi-Helm *et al*., 2019; Ing-Simmons *et al*., 2021). We speculate that Mod(mdg4)-N activity in chromatin boundary formation supports the developmental program (Fig. 6B) as a severe ovulation defect is observed upon Mod(mdg4)-N loss (Fig. 1G-I). Also, we suggest that there is functional redundancy among insulator proteins, and it is plausible that other insulator proteins compensate for loss of Mod(mdg4)-N, because not all Mod(mdg4)-N binding sites show Pol II loss or TAD insulation loss upon Mod(mdg4)-N depletion (Fig. 4,5). Similarly, loss of Beaf-32, which is the most enriched insulator protein at TAD boundaries, results in only moderate changes in chromatin conformation (Ramírez *et al*., 2018), indicating that functional overlaps among insulator proteins are more common than previously thought.

In summary, both mechanisms of Mod(mdg4)-N are likely to play important roles in orchestrating proper gene expression, and further functional dissections of insulator proteins at the molecular level will lead us to understand why TAD disruption causes minor effects although mutations of insulator proteins show a drastic effect on phenotypes.

## DATA AND CODE AVAILABILITY

GSE (Micro-C XL, ChIP-Seq): GSE176196 Github (python Script): https://github.com/Chikara-Takeuchi/2021-Mod-mdg4

## Acknowledgements

We thank all Siomi lab members for experimental support and fruitful discussions, especially Akihiko Sakashita and Hirotsugu Ishizu for critical reading of the manuscript. Pol II antibody used in this study was a kind gift from Hiroshi Kimura. CP190 antibody used in this study was a kind gift from Elissa Lei. siRNAs of each Mod(mdg4) variant were designed by Hidetoshi Tahara (Advanced Animal Model Support platform).

CT is supported by funding from JST JPMJSP2123 and Koyo Sakaguchi Memorial Keio University Medical Science Fund. YWI is supported by funding from JSPS KAKENHI Grant Numbers 19H05268 and 18H02421, JST PRESTO Grant Number JPMJPR20E2. HS is supported by funding from JSPS KAKENHI Grant Number JP 16H06276, 19H05753. T.F. is supported by the Grant-in-Aid for Transformative Research Areas (A) (Research in a Proposed Research Area) (21H05742), the Grant-in-Aid for Scientific Research on Innovative Areas (Research in a Proposed Research Area) (20H05357), the Grant-in-Aid for Scientific Research (B) (19H03154) from the Japan Society for the Promotion of Science, research grants from the Takeda Science Foundation, the Sumitomo Foundation, the Senri Life Science Foundation and the Mitsubishi Foundation. M.Y is supported by the Grant-in-Aid for Early-Career Scientists (20K15710) from the Japan Society for the Promotion of Science and JST, ACT-X (JPMJAX211J). We thank Katrina Woolcock (Life Science Editors) for editing services.

## Author Contributions

CT designed and performed most of the experiments and analyzed the data. MY and TF performed live-imaging for the enhancer blocking assay in embryo. AS performed immunofluorescence experiment and fly fertility assay. YWI established antibody against Mod(mdg4) common region and performed ChIP-Seq for CP190. SK and KS generated transgenic Mod(mdg4)-N mutant fly. YWI, HS conceived this study. CT, YWI, HS wrote the paper with input from the other authors.

## Materials and Methods

## KEY RESOURCE TABLE

See additional file

## LEAD CONTACT AND MATERIALS AVAILABILITY

Further information and requests for resources and reagents should be directed to and will be fulfilled by the Lead Contact, Yuka W Iwasaki.

## EXPERIMENTAL MODEL AND SUBJECT DETAILS

### Cell lines and culture condition

Ovarian somatic cells (OSC) were cultured as described (Niki et al., 2006; Saito *et al*., 2009). Briefly, OSC were cultured in Shield and Sang M3 Insect Medium (Sigma) supplemented with 10% fly extract, 10% fetal bovine serum, 0.6 mg/ml glutathione, and 10 μg/ml insulin. OSC were passed every second day.

### Fly strains

New alleles of mod(mdg4) were generated using the transgenic Cas9 system as previously described (Kondo and Ueda, 2013). The 20-bp sequence of the gRNA targeting the RN variant-specific exon of mod(mdg4) is as follows: GTAGTTGCGGAACACCAGCT. Multiple candidate mutant lines were established from individual male offspring of parents carrying both the nos-Cas9 and U6-gRNA transgenes. PCR was performed on their genomic DNA to amplify a region surrounding the gRNA target and their sequences were determined to search for indel mutations. Two lines, RN1-2 and RN1-4, were found to carry distinct frameshift mutations (Fig. S2A) and were subjected to further functional analysis.

In all live-imaging experiments, *Drosophila melanogaster* embryos at nuclear cycle 14 were analyzed. The following fly lines were used in this study: *nanos>MCP-GFP, His2Av-mRFP/CyO* (Yokoshi et al., 2020), *DSCP*_*WT*_*-MS2-yellow-sna shadow enhancer* (Yokoshi *et al*., 2020), *DSCP*_*WT*_*-MS2-yellow-gypsy-sna shadow enhancer* (Yokoshi et al., 2021), *DSCP*_*WT*_*-MS2-yellow-HeT-A-sna shadow enhancer* (this study), *DSCP*_*WT*_*-MS2-yellow-HETRP-sna shadow enhancer* (this study), *DSCP*_*WT*_*-MS2-yellow-HeT-A-sna shadow enhancer*-*HeT-A* (this study), *DSCP*_*WT*_*-MS2-yellow-HETRP-sna shadow enhancer-HETRP* (this study).

## METHOD DETAILS

### Cell transfection

For transfection of small interfering (si)RNA, 200 pmol siRNA duplex was transfected using the Cell Line 96-well Nucleofector Kit SF (Lonza) and program DG150 of the 96-well Shuttle Device (Lonza). For transfection of expression vectors, expression vectors were transfected using Xfect Transfection Reagent (TaKaRa Clontech), in accordance with the manufacturer’s instructions. All siRNA sequences used in this study are listed in Supplementary Table 1.

### Generation of antibodies

The 1469-1901 bp region of Mod(mdg4) variant N mRNA (Flybase: FBtr0084073) and the 3984-4511 bp region of HeT-A 23Zn-1(GenBank: U06920.2) were subcloned into pMAL-c2G and pGEX-5X-1 from OSC or ovary cDNA. These vectors were expressed in Rosseta-gami B(DE3) (Novagen) with 1 mM IPTG at 18°C overnight. MBP-tagged proteins and GST-tagged protein were purified with Amylose Resin (New England Biolabs) and Glutathione Sepharose 4B (Cytiva). These purified proteins were used as antigens, and hybridomas with SP2/O were generated and screened according to standard protocol (Greenfield, 2018).

### Quantitative reverse-transcription PCR

For OSC, RNA extraction and cDNA preparation were performed with SuperPrep II Cell Lysis & RT kit for qPCR from 1.0 × 10^5^ OSC. For fly ovaries, total RNA extraction was performed with ISOGEN II (Nippon Gene: 311-07361) and cDNA was synthesized with Transcription First Strand cDNA Synthesis Kit (Roche 04379012001). TB Green Premix Ex Taq II (Clontech) was used for quantitative PCR with indicated qPCR primers (Supplementary Table 1).

### Western blot

Samples were lysed to the concentration of 5.0 × 10^5^ OSC / 1 ovary per 10 µl 1× Laemmli buffer and heated at 95°C for 5 min. Proteins were separated by SDS–polyacrylamide gel electrophoresis (PAGE) and transferred to a 0.45 µm nitrocellulose membrane (Cytiva). Membrane was washed by PBS and blocked with 3% skim milk in PBS-T, then incubated with primary antibody for 1 h at room temperature. For primary antibody, anti-beta tubulin (1:5000), anti-c-Myc (1:1000), anti-Ty1 antibody (1:2000), anti-HeT-A Gag supernatant (1:1), anti-Mod(mdg4) common region supernatant (1:4) and anti-Mod(mdg4) variant N supernatant (1:1) were used with indicated dilutions. After three washes with PBS-T, membrane was incubated with HRP-conjugated secondary antibody (1:5000) for 30 min, followed by three PBS-T washes. The membranes were incubated at room temperature for 1 h. The membrane was incubated with ECL Prime Western Blotting Detection Regent (Cytiva) and was exposed to Amersham Hyperfilm ECL (Cytiva). Exposed film was developed by X-ray film developer (KONICAMINOLTA TCX-101).

### Immunofluorescence

Ovaries were fixed with 4% paraformaldehyde for 20 min and washed once by PBS-T (0.2% Tween20 in PBS), followed by blocking by PBS-BT (0.2% Tween20 and 10 mg/ml BSA in PBS). Primary antibodies were diluted with PBS-BT and incubated with samples overnight at 4°C. For primary antibodies, anti-Orb (1:200) and anti-Vasa supernatant (1:1) (Sumiyoshi et al., 2016) were used with indicated dilutions.

Following three washes with PBS-T, secondary fluorophore-conjugated antibody was diluted at 1:1000 in PBS-BT and incubated with samples for 2 h at room temperature. The stained samples were washed three times with PBS-T and mounted by DAPI-containing medium. Samples were imaged with Zeiss LSM-710 confocal microscope.

### Isolation of Ty1-Mod(mdg4) variants stable line OSC

Stable cell lines were established as described previously (Takeuchi et al., in press). In short, Mod(mdg4) variants were cloned into pPB-2×Ty1-Tjen-EGFP-P2A-BlastR. Stable clones were selected by medium containing 50 µg/ml blasticidin.

### Plasmid construction

#### HeT-A/HETRP sequence cloning

HeT-A or HETRP sequences were cloned from the OSC genome. These sequences were amplified by PCR using specific primers (rep_HeT-A_F/R or rep_HETRP_F/R), and cloned to pBlueScript SK(+) EcoRI/NotI treated fragment by using NEBuilder HiFi DNA Assembly Master Mix (NEB). These plasmids are named pBS-HeT-A or pBS-HETRP respectively. Primer sequences for cloning are shown in Supplementary Table 1.

#### DSCP_WT_-MS2-yellow-HeT-A-sna shadow enhancer

A DNA fragment containing HeT-A sequence was amplified from pBS-HeT-A using primers (5’
s-ACA TGA AGC TTC TTC TCC GTT CTA CCT CAA T-3’) and (5’-ACG GGA AGC TTC ACA GGG TGC CGC AAA AAT TG-3’) and digested with HindIII. The resulting fragment was inserted into the unique HindIII site of pbphi-DSCP-MS2-yellow*-*sna shadow enhancer (Yokoshi *et al*., 2020).

#### DSCP_WT_-MS2-yellow-HETRP-sna shadow enhancer

A DNA fragment containing HETRP sequence was amplified from pBS-HETRP using primers (5’
s-ACG GGA AGC TTG TGT GTC ATC CAT TTC GTT T-3’) and (5’-ACA TGA AGC TTC GAC GCG TAC ACA TAT TTC G -3’) and digested with HindIII. The resulting fragment was inserted into the unique HindIII site of pbphi-DSCP-MS2-yellow*-*sna shadow enhancer (Yokoshi *et al*., 2020).

#### DSCP_WT_-MS2-yellow-HeT-A-sna shadow enhancer-HeT-A

A DNA fragment containing HeT-A sequence was amplified from pBS-HeT-A using primers (5’
s-ACA TGG CTA GCC TTC TCC GTT CTA CCT CAA TAT ATC-3’) and (5’-TTA AAG CTA GCC ACA GGG TGC CGC AAA AAT TG-3’) and digested with NheI. The resulting fragment was inserted into the unique NheI site of *DSCP*_*WT*_*-MS2-yellow-HeT-A-sna shadow enhancer*.

#### DSCP_WT_-MS2-yellow-HETRP-sna shadow enhancer-HETRP

A DNA fragment containing HeT-A sequence was amplified from pBS-HETRP using primers (5’
s-ACA TGG CTA GCG TGT GTC ATC CAT TTC GTT TAT TC-3’) and (5’-TTA AAG CTA GCC GAC GCG TAC ACA TAT TTC GC -3’) and digested with NheI. The resulting fragment was inserted into the unique NheI site of *DSCP*_*WT*_*-MS2-yellow-HETRP-sna shadow enhancer*.

### Site-specific transgenesis by phiC31 system

All reporter plasmids were integrated into a unique landing site on the third chromosome using VK00033 strain (Venken et al., 2006). PhiC31 was maternally provided using *vas-phiC31* strain (Bischof et al., 2007). Microinjection was performed as previously described (Ringrose, 2009). In brief, 0-1 h embryos were collected and dechorionated with bleach. Aligned embryos were dried with silica gel for ∼7 min and covered with FL-100-1000CS silicone oil (Shin-Etsu Silicone). Subsequently, microinjection was performed using FemtoJet (Eppendorf) and DM IL LED inverted microscope (Leica) equipped with M-152 Micromanipulator (Narishige). Injection mixture typically contains ∼500 ng/μl plasmid DNA, 5 mM KCl, 0.1 mM phosphate buffer, pH 6.8. The mini-white marker was used for screening.

### MS2 live-imaging

Virgin females of *nanos>MCP-GFP, His2Av-mRFP/CyO* (Yokoshi *et al*., 2020) were mated with males carrying the MS2 allele. The resulting embryos were dechorionated and mounted between a polyethylene membrane (Ube Film) and a coverslip (18 mm x 18 mm), and embedded in FL-100-450CS (Shin-Etsu Silicone). Embryos were imaged using LSM900 (Zeiss). Temperature was kept between 23.5 and 24.5°C during imaging. Plan-Apochromat 40x / 1.4 N.A. oil immersion objective was used. A stack of 26 images separated by 0.5 μm was acquired at each time point, and the final time resolution was 16.8 sec/frame. Images were captured in 16-bit. Images were typically taken from the end of nuclear cycle 13 to the onset of gastrulation at nuclear cycle 14. During imaging, data acquisition was occasionally stopped for a few seconds to correct z-position, and data were concatenated afterwards. For each cross, three biological replicates were taken. The same laser power and microscope setting were used for each set of experiments. The laser power was measured using X-Cite XR2100 power meter (Lumen Dynamics).

### Chromatin immunoprecipitation (ChIP)

ChIP experiments of Ty1-tagged Mod(mdg4) variants and CP190 (Moshkovich et al., 2011)were performed using the truChIP Chromatin Shearing Kit according to the manufacturer’s instructions with minor modifications. 2 × 10^7^ OSC were fixed with 1% formaldehyde for 10 min and quenched and lysed with buffers from the truChIP Chromatin Shearing Kit. Fixed chromatin was sheered with Bioruptor II (BMbio BR2012A) for 20 cycles of 30 s ON/30 s OFF, high settings. Sheered chromatin was immunoprecipitated by antibody-conjugated Dynabeads protein G for 2 h. Immunoprecipitated chromatin was incubated with Proteinase K and RNase, and de-crosslinked at 65°C overnight.

For ChIP experiments of RNA polymerase II, 2 × 10^7^ OSC were fixed with 1% formaldehyde for 10 min and quenched with final 0.1 M Glycine for 5 min. After washing twice with PBS, nuclei were isolated with 1 ml of swelling buffer (25 mM HEPES-KOH (pH 7.5), 1.5 mM MgCl_2_, 10 mM KCl, 0.1% NP-40, 1 mM DTT and 1x Protease Inhibitor). Isolated nuclei were lysed in 400 µl of sonication buffer (50 mM HEPES-KOH (pH 7.4), 140 mM NaCl, 1 mM EDTA, 1% Triton X-100, 0.1% Na-deoxycholate, 0.1% SDS and 1× Protease Inhibitor) and were sheered with Bioruptor II (BMbio BR2012A) for 20 cycles of 30 s ON/30 s OFF, high settings. 3 µg of Pol II antibody were incubated with chromatin overnight (Stasevich et al., 2014). The next day, 20 µl of Dynabeads M-280 Sheep anti-Mouse IgG was added to the sample and incubated for 1 h. Beads were washed and treated with Proteinase K and RNase as described (Haring et al., 2007). DNA was purified with isopropanol precipitation using Pellet Paint NF Co-precipitant (Merck: 70748).

Fragments from the ChIP experiment were sheared to ∼200 bases using Covaris S220. These were used for library preparation with the NEBNext Ultra II DNA Library Prep Kit for Illumina (NEB) following the manufacturer’s protocol.

### Micro-C XL

Micro-C XL was performed as previously described (Hsieh *et al*., 2020). 1.5 × 10^7^ OSC were fixed with 1% formaldehyde for 10 min and quenched with final 0.75 M Tris-HCl (pH 7.5) for 5 min. After washing twice, OSC were fixed again with final 3 mM DSG for 45 min at room temperature and quenched with final 0.75 M Tris-HCl (pH 7.5) for 5 min. Nuclei were isolated with 100 µl MBuffer#1 (50 mM NaCl, 10 mM Tris-HCl pH 7.5, 5 mM MgCl_2_, 1 mM CaCl_2_, 0.2% NP-40 and 1× Protease inhibitor (EDTA free)) and were incubated on ice for 20 min. After washing with MBufer#1, nuclei were resuspended in MBuffer#1 again and 4,000 gel units of MNase were added to solution, followed by incubation at 37°C for 10 min with rotation. Reaction was stopped by adding 1 µl of 0.5 M EGTA (pH 8.0) and nuclei were washed twice with MBuffer#2(50 mM NaCl, 10 mM Tris-HCl pH 7.5, 10 mM MgCl_2_). DNA ends were phosphorylated by 45 µl end-chewing mix (5 µl 10×NEBuffer 2.1, 10 µl ATP (10 mM), 2.5 µl DTT (100 mM), 2.5 µl T4PNK (10 U/µl), 25 µl Milli-Q) and incubated at 37°C for 15 min. After incubation, 5 µl of Klenow fragment (5 U/µl) was added and incubated at 37°C for 15 min again. DNA overhangs were filled with biotin by adding 25 µl end-labeling mix (5 µl biotin-14-dATP (1 mM), 5 µl biotin-14-dCTP (1mM), 0.5 µl d[G+T]TP(10 mM), 2.5 µl 10× T4 DNA Ligase Buffer, 0.25 µl BSA (10 mg/ml), 11.75 µl Milli-Q), and incubated at 25°C for 45 min. End labeling was stopped by adding 5 µl 0.5 M EDTA and inactivated by incubation at 65°C for 20 min. Collecting by centrifugation (12000 *g* for 5 min), samples were washed with 1 ml MBuffer#3 (50 mM Tris-HCl pH7.5, 10 mM MgCl_2_). Chromatin pellet was resuspended in 250 µl End-ligation mix (25 µl 10× T4 DNA Ligase Buffer, 2.5 µl BSA (10mg/ml), 12.5µl T4 DNA ligase (400 U/µl), 210 µl Milli-Q) and rotated slowly at room temperature for 4 h. Collecting by centrifugation (16000 *g* 5 min), pellet was resuspended in Exonuclease solution (10 µl 10× NEBuffer #1, 5 µl Exonuclease III (100 U/µl), 85 µl Milli-Q) and sample was incubated at 37°C for 15 min to remove biotin from unligated ends. For deproteination and reverse crosslinking, 25 μl Proteinase K (20 mg/ml), 15 µl RNase (0.5 mg/ml) and 15 µl 10% SDS were added and the sample was incubated at 65°C o/n. Ligated DNA was purified with isopropanol precipitation using Pellet Paint NF Co-precipitant (Merck) and resuspended in 150 µl Milli-Q. 5 μl Dynabeads MyOne Streptavidin C1 were washed twice with 300 μl Tween wash buffer (5 mM Tris-HCl, pH 7.5, 0.5 mM EDTA, 1 M NaCl) and suspended in 150 μl 2× Biotin binding buffer (10 mM Tris-HCl, pH 7.5, 1 mM EDTA, 2 M NaCl). 150 μl washed Streptavidin beads in 2× TBW were added to the sample and incubated with rotation at RT for 20 min. Beads were washed twice with 300 μl Tween wash buffer. These were used for library preparation with the GenTrack library preparation kit following the manufacturer’s protocol. To remove monomer or tetramer sized fragments, size-selection was performed with SPRI select after amplification of library.

### Enhancer blocking reporter assay in OSC

Reporters were constructed from OSC_Reporter_UAS_traffic jam_BoxB_d2eGFP_t2a_Blast (Batki et al., 2019) and pAc5.1 vector in which Flag-mCherry:P2A:PuroR gene was cloned using the indicated primer to add an additional restriction site (Supplemental Table 1). Wild-type and mutant *HETRP* sequences were inserted into reporter with NruI or SalI restriction enzyme and NEBuilder hifi DNA assembly (NEB). These reporters were transfected with pHsp70-Myc-PBase plasmid as described (Takeuchi *et al*., in press). Cells were selected with puromycin and fluorescence was measured with SH800Z (Sony). Normalized EGFP intensity is original EGFP signal intensity divided by mCherry intensity.

### Fertility assay

7-10 virgin female flies were collected 3-5 days before mating. Virgin female flies were mated with the indicated male and kept at 25°C overnight. The next day, mated flies were transferred onto grape juice plate and kept at 25°C without light. After 3 h, eggs were counted and kept at 25°C again. After 24 h, hatching eggs were counted.

## QUANTIFICATION AND STATISTICAL ANALYSIS

### Image analysis

All the image processing methods and analysis were implemented in MATLAB (R2020a, MathWorks).

### Segmentation of nuclei

For each time point, maximum projection was obtained for all z-sections per image. His2Av-mRFP was used to segment nuclei. 512 × 512 maximum projection images were initially cropped to 300 × 430 to remove nuclei at the edge, and used for subsequent analysis. For nuclei segmentation, His2Av images were first blurred with Gaussian filter to generate smooth images. Pixels expressing intensity higher than 5% of the global maxima in the histogram of His2Av channel were removed. Processed images were converted into binary images using a custom threshold-adaptative segmentation algorithm. Threshold values were determined at each time frame by taking account of (i) histogram distribution of His2Av channel and (ii) the number and the size of resulting connected components. Boundaries of components were then modified to locate MS2 transcription dots inside of nearest nuclei. In brief, pixels with intensity twice larger than mean intensity of MS2 channel were considered as transcription dots, and new binary image was created for each time frame. The Euclidean distances between the centroid of binarized transcription dot and all boundaries of segmented nuclei were calculated. Boundary of the nucleus with the smallest Euclidean distance was modified in order to capture transcription dot within a nucleus. Centroids of connected components in nuclei segmentation channel were used to compute the Voronoi cells of the image. Resulting binary images were manually corrected using Fiji (https://fiji.sc).

### Tracking of nuclei

Nuclei tracking was done by finding the object with minimal movement across the frames of interest. For each nucleus in a given frame, Euclidean distances between the centroids of the nucleus in the current time frame and the nuclei in the previous time frame were determined. Nucleus with the minimum Euclidean distance was considered as a same lineage.

### Recording of MS2 signals

3D raw images with all z-sections of MCP-GFP channel were used to record MS2 fluorescence signals. Using segmented regions from max projected images of His2Av-mRFP channel, fluorescence intensities within each nucleus were extracted. 3D fluorescence values were assigned to the nearest segmented regions of projected images. Signals of MS2 transcription dots were determined by calculating an integral of fluorescence intensities around the brightest pixel within each nucleus using a 2D Gaussian fitting method as described below. (i) The xyz position of transcription site was determined as the brightest pixel in each nucleus. (ii) A 2D Gaussian fitting was performed in a 11 × 11 pixels region with a single z-plane centering the transcription site to estimate a fluorescent dot intensity and a local background. Fitting was performed with the following formula

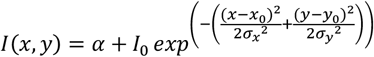

where *α* is the local background intensity, *I*_*0*_ is the amplitude of the peak fluorescence intensity, *x*_*0*_ and *y*_*0*_ are the center of the peak, *σ*_*x*_ and *σ*_*y*_ are the spreads of the fluorescent dot. (iii) The intensity of MS2 transcription dot was calculated as 2*πσ*_*x*_*σ,I*_*y*_ from fitting parameters as an estimated integral value after subtracting the local background (Martin et al., 2013). Subsequently, minimum MS2 intensities were determined for individual trajectories and subtracted to make the baseline zero.

### Detection of transcriptional bursting

A transcriptional burst was defined as a local change in fluorescence intensity. First, signal trajectories were smoothed by averaging within a window of 5 timeframes. When a nucleus had above-threshold transcription activity, burst was considered to be started. Burst was considered to be ended when the intensity dropped below 55% of the local peak value. When the burst duration was less than 5 timeframes, it was considered as a false-positive derived from detection noise. When the signal trace exhibited continuous decreasing at the beginning of burst detection, it was also not considered as a burst. Location of defined burst was then moved two timeframes afterwards to better capture the center of individual bursting event. The same method and threshold value were used for each set of experiments.

### Description of bursting properties

From each trajectory, number of bursts, amplitude and duration of each burst, and total integrated signal (output) produced by each nucleus were measured. To determine amplitude, the peak value during the burst was measured using trajectories after smoothing by averaging within a window of 5 timeframes. Duration was determined by measuring the length of each burst. Total RNA production was measured by taking the area under the raw trajectory. Amplitude and duration for each nucleus were determined by taking the average of all analyzed bursts in a single nucleus.

### mRNA-Seq data analysis

Reads were mapped to dm6 by STAR (Dobin et al., 2013) using the following parameter (STAR --runThreadN 12 –genomeDir {path_to_index} --readFilesIn rawdata/RNA_egfp.fastq -- outSAMtype BAM SortedByCoordinate -- outFilterMultimapNmax 100 --chimOutType WithinBAM -- chimSegmentMin 20 --genomeLoad NoSharedMemory). Then, reads mapped to transposable elements were counted by featureCounts (Liao et al., 2014)using the following parameter (-M --fraction -F GTF -t exon) with annotation from repeatmasker (https://www.repeatmasker.org/) (Teissandier et al., 2019).

### ChIP-Seq data analysis – mapping and peak call

Adaptors added by NEBNext Ultra II were cut by cutadapt (Martin, 2011)using the following parameters (cutadapt -j 12 -a AGATCGGAAGAGCACACGTCTGAACTCCAGTCAC -A AGATCGGAAGAGCGTCGTGTAGGGAAAGAGTGTAGATCTCGGTGGTCGCCGTATCATT -m 20 -o trim/${name2}_R1_trim.fastq.gz -p trim/${name2}_R2_trim.fastq.gz raw_file/${name2}_R1.fastq.gz raw_file/${name2}_R2.fastq.gz). For mapping to the genome, subsequent reads were mapped to the dm6 by bowtie2 using the following parameters (bowtie2 -p 16 -x ${reference}/dm6 -N 1 - 1 trim/${name2}_R1_trim.fastq.gz -2 trim/${name2}_R2_trim.fastq.gz -S sam_file/${name2}.sam). Peak call was performed by MACS using the following parameter: for Mod(mdg4) variants(macs2 callpeak -t merge_file/Ty-AF_merge.sort.bam -c bam_file/Input-AF.sort.bam -n Ty-AF -f BAM -q 1e-15 -g dm --outdir peakcall_strict) and for PolII(macs2 callpeak -t ${item} - c ../bam_file/Input_All.sort.bam -n ${item%%.bam} -f BAM -g dm --outdir ../peakcall). To note, peak call for Mod(mdg4) was performed with strict cutoff values to rule out ambiguous peaks and we confirmed almost no peak was detected in Ty1-tag ChIP for wildtype OSC dataset. Binding motif was found with MEME-ChIP (Machanick and Bailey, 2011)with standard options. Coverage tracks were visualized by pyGenomeTracks (Lopez-Delisle et al., 2020) with standard options.

### ChIP-Seq data analysis – handling of Pol II ChIP-Seq data

Pausing index was calculated as previously described (Gilchrist *et al*., 2010). Pol II signal of both promoter region (TSS-250 ∼ TSS+250) and gene body (TSS+500∼TES-500) was calculated from a public dataset (Sienski *et al*., 2012). Genes whose length is shorter than 1500 bp were removed from downstream analysis. To obtain the pausing index for each gene, we divided reads of promoter region by reads of gene body. Cumulative distribution plot was illustrated by Python. For identifying Pol II loss genes, maximum coverage was calculated for each Pol II peak. Fold change of this maximum value was calculated between EGFP KD and Mod(mdg4) variant N KD.

### Micro-C XL data analysis - General processing

Distiller (https://github.com/open2c/distiller-nf) pipeline was used to process Micro-C datasets. First, sequencing reads were mapped to dm6 using bwa mem with flags-SP. Second, mapped reads were parsed and classified using the pairtools package (https://github.com/open2c/pairtools) with “--walks-policy all” option to get cool file. PCR/optical duplicates are removed with 1bp flexibility. Pairs were filtered using mapping quality scores (MAPQ > 30) on each side of aligned chimeric read, binned into multiple resolutions (200 bp to 50k bp) and low coverage bins were removed. Finally multiresolution cooler files were created using the cooler package (https://github.com/open2c/cooler.git) (Abdennur and Mirny, 2019). We normalized contact matrices using the iterative correction procedure (Imakaev et al., 2012). All browser snapshots of Micro-C contact map were generated by the HiGlass browser unless otherwise mentioned (Kerpedjiev et al., 2018).

### Micro-C XL data analysis – P(s) curve

P(s) curve was calculated as previously described (Naumova *et al*., 2013) using cooltools. In short, we first divided all genomic separations into logarithmically spaced 50 bins between 500 bp and 10 Mbp. Reads mapping to chr2R, 2L, 3R, 3L, 4 and X were used for calculation. To obtain P(s), we divided the number of reads in each bin by the number of possible fragment pairs and normalized P(s) such that the integral of P(s) over the range of distances was 1.

### Micro-C XL data – Boundary analysis

Diamond insulation score (Crane et al., 2015) was calculated by using cooltools (https://github.com/open2c/cooltools) at each resolution with (resolution(bp)×25) sized window. Regions with local minima are defined as boundaries.

For downstream analysis, we used only “strong boundary” proposed in previous research to avoid ambiguous boundaries (Akgol Oksuz et al., 2021). In short, we set cut-off of boundary strength to mean of all called boundary strength (weak boundary < mean < strong boundary). Overlapping between peaks of each ChIP-Seq and boundaries was calculated with bedtools intersect. To identify differentially insulated region, we compared insulation score of all bins between EGFP KD and Mod(mdg4) variant N KD at 200 bp resolution. Bins with low coverage region (i.e., TE rich region or satellite repeat) were excluded from downstream analysis. Differentially insulated regions were defined as top or bottom 0.1% at fold change of insulating score. Adjacent bins were merged with bedtools merge with -d 250 options. Metaplots were illustrated using the plotHeatmap command in deeptools.

## KEY RESOURCES TABLE

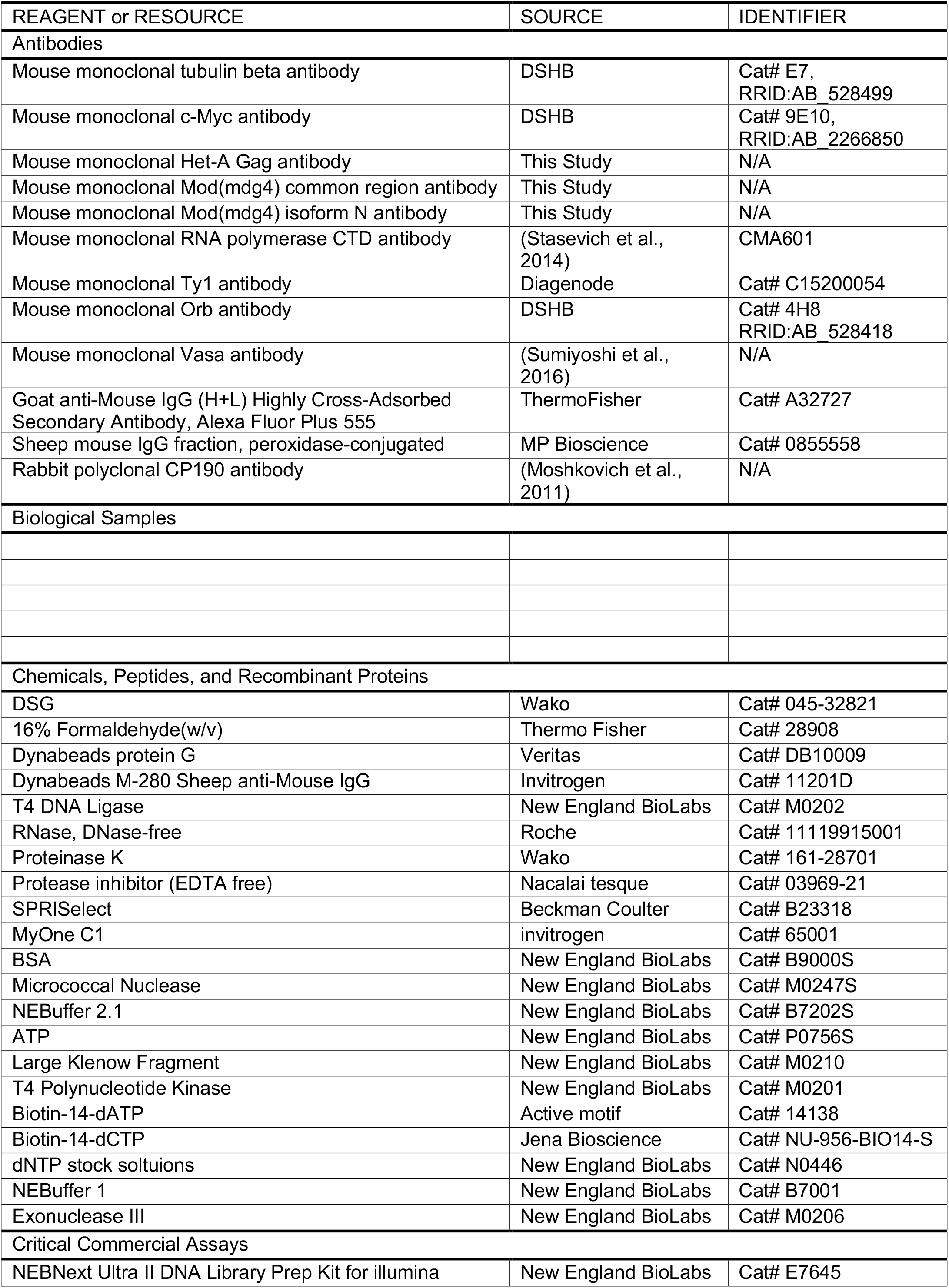
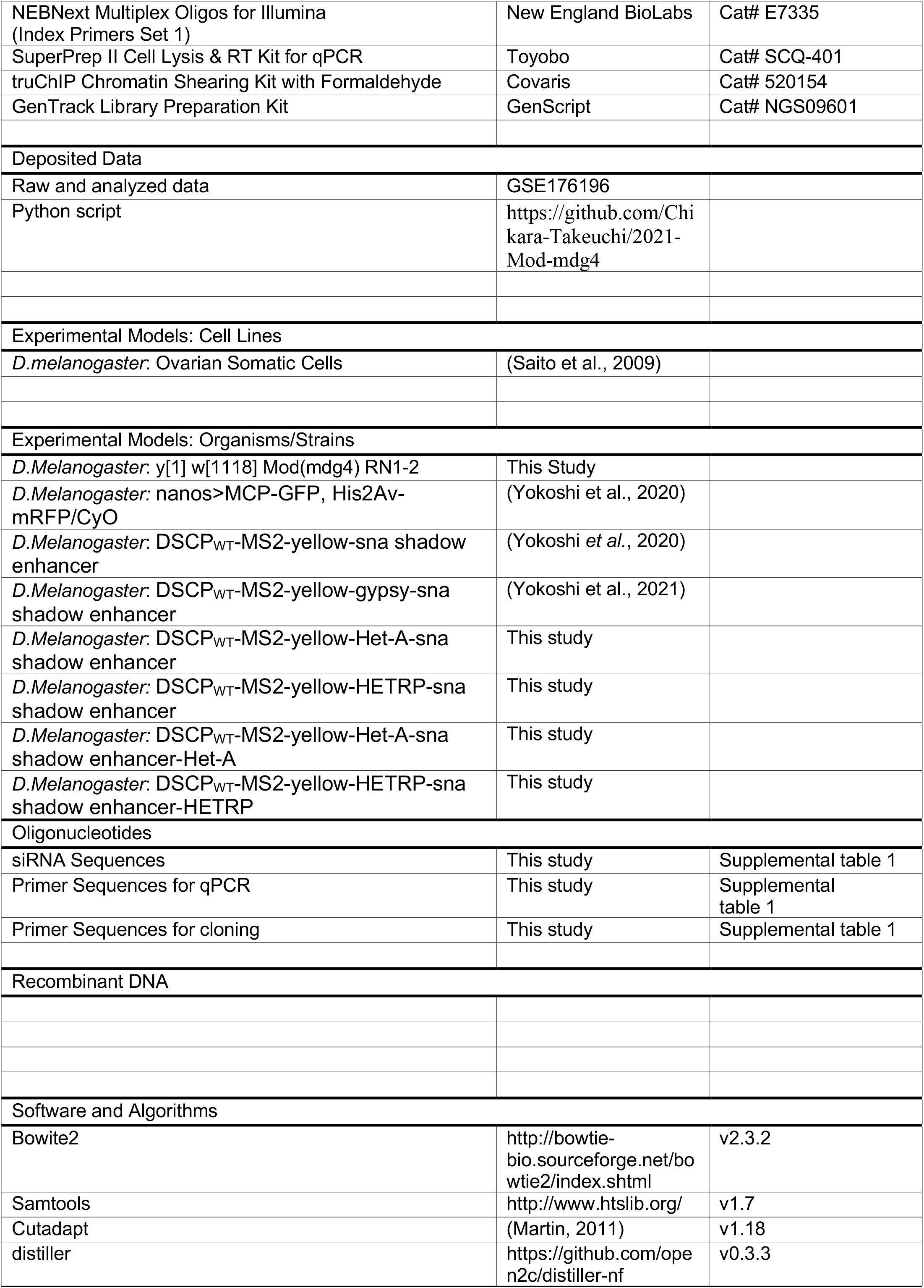
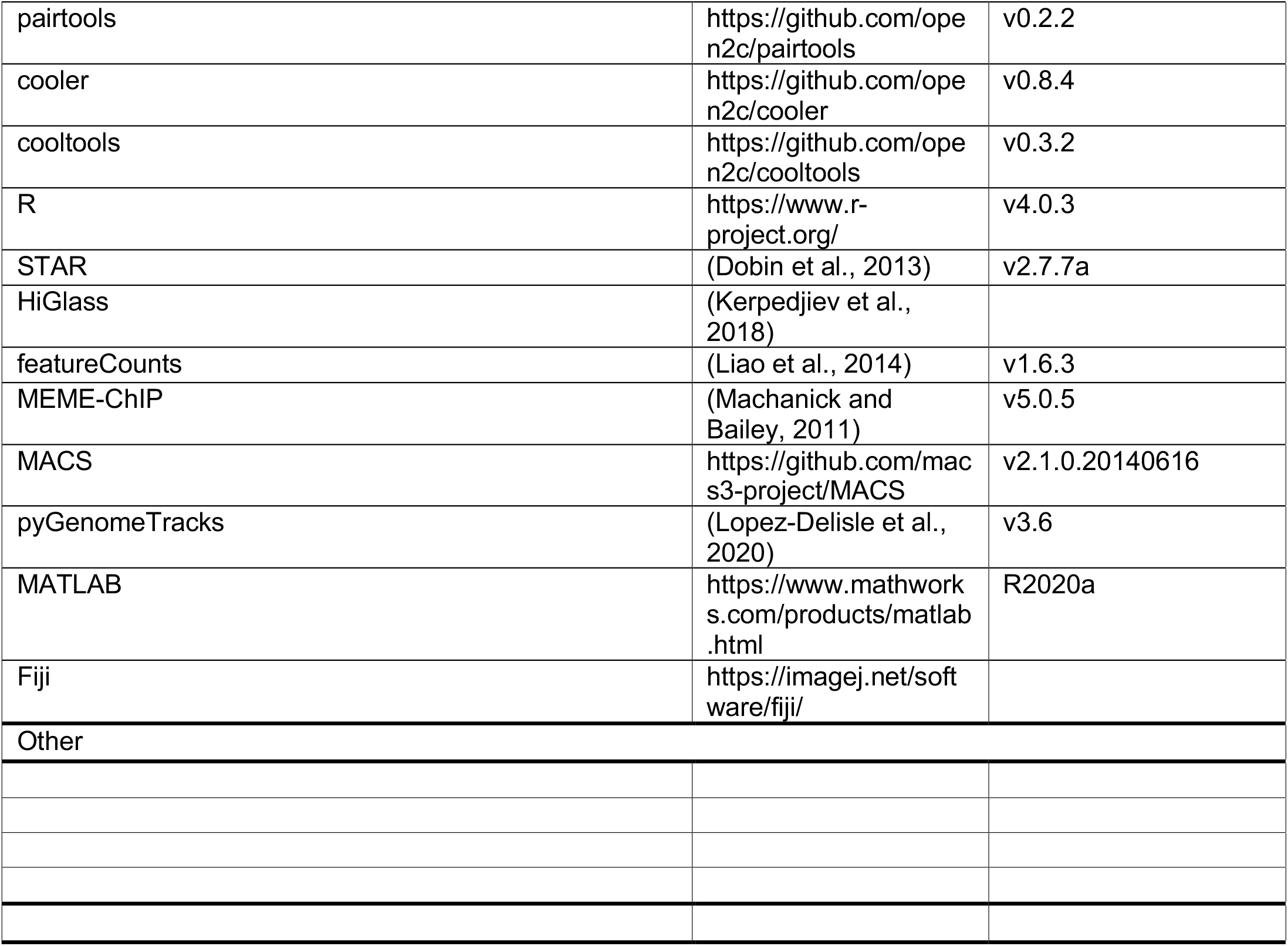

**Fig. S1.**
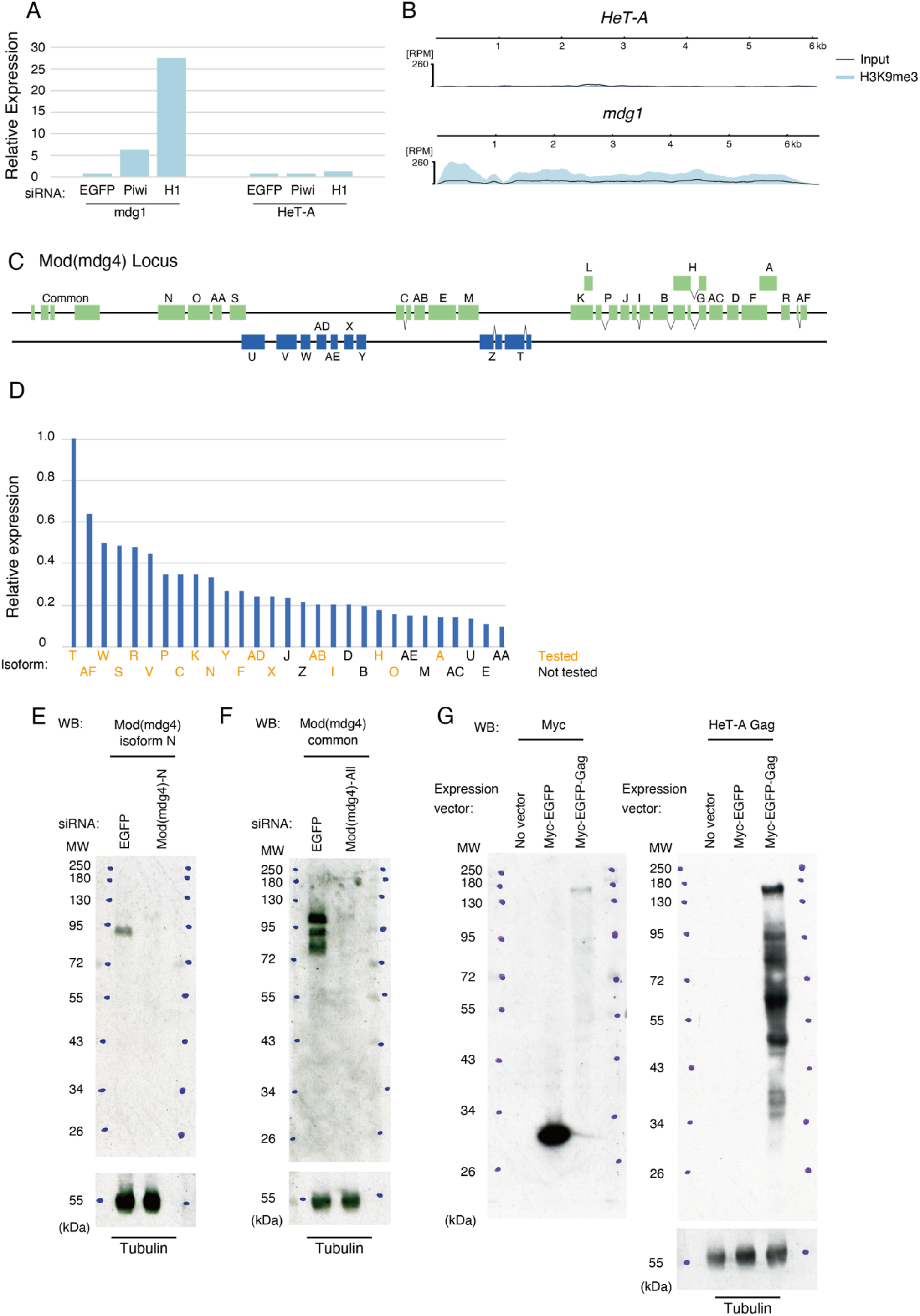
related to Fig. 1. **A**, Relative expression calculated from RNA-Seq reads mapped to *mdg1* and *HeT-A* upon EGFP (control), piwi, or H1 KD. Read counts are normalized by reads per million (RPM) and the expression level at EGFP KD as 1. **B**, Coverage tracks showing results of H3K9me3 ChIP-Seq (blue) and its input (black line) mapped onto the consensus sequence of *mdg1* and *HeT-A* from repbase. Coverages are normalized by counts per mapped million reads. **C**, Schematic diagram of Mod(mdg4) locus. Mod(mdg4) has 4 common exons and 31 variant-specific exons. The names of variants are indicated on the exons. **D**, Bar plot showing relative expression of each Mod(mdg4) variant in OSC as revealed by EGFP-KD RNA-Seq. Variants G and L have no unique sequence and are therefore excluded. The names of variants tested in the siRNA KD screen (Fig. 1D) are indicated with orange. **E-G**, Western blotting shows the specificity of the antibodies used in this study. Antibodies against Mod(mdg4) variant N (E), Mod(mdg4) common region (F) and HeT-A Gag (G) were used. We checked the specificity of antibody by KD of endogenous Mod(mdg4) in OSC (E,F). Because HeT-A Gag protein is not expressed in normal conditions, we checked the specificity of antibody by comparing no vector control, Myc-EGFP or Myc-EGFP-Gag protein overexpression (G) in OSC. Tubulin is detected as a loading control for each experiment. Molecular weight (MW) is indicated at the left of each image.

**Fig. S2.**
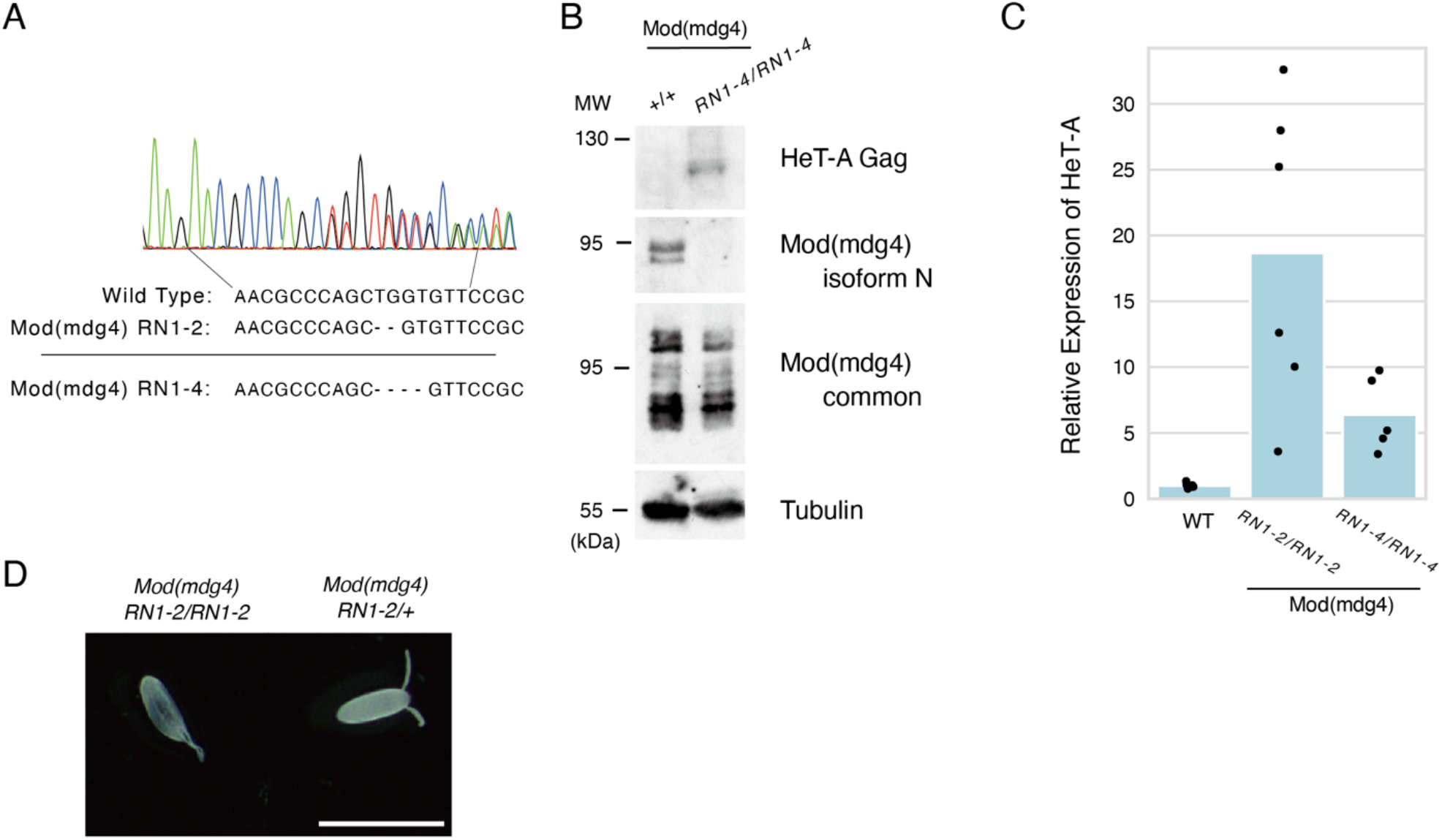
Analysis of Mod(mdg4)-N mutants. **A**, Fly strain (*Mod(mdg4) RN1-2* and *RN1-4*) used in this study. The result of sanger sequencing of *Mod(mdg4) RN1-2* heterozygote is shown in upper panel. Their sequences are indicated below. *Mod(mdg4) RN1-2* and *RN1-4* have two or four base pairs missing from their variant-specific exon. **B**, WB showing protein levels of HeT-A Gag, Mod(mdg4)-N, total Mod(mdg4) and tubulin (loading control) in ovaries of *+/+* wildtype (OregonR) or homozygous *Mod(mdg4) RN1-4/RN1-4* mutant. Molecular weight (MW) is indicated at the left of each image. **C**, qRT-PCR showing RNA expression of *HeT-A*. y-axis is normalized by RP49 (n=6 for WT or *Mod(mdg4) RN1-2/RN1-2* and n=5 for *Mod(mdg4) RN1-4/RN1-4*). Average of WT ovary is normalized as 1. Each dot indicates values obtained by different experiments. **D**, Image of eggs ovulated from Mod(mdg4) RN1-2/1-2 homozygous mutant (left) and heterozygous mutant (right) obtained under an inverted microscope. White bar indicates 1 mm.

**Fig S3.**
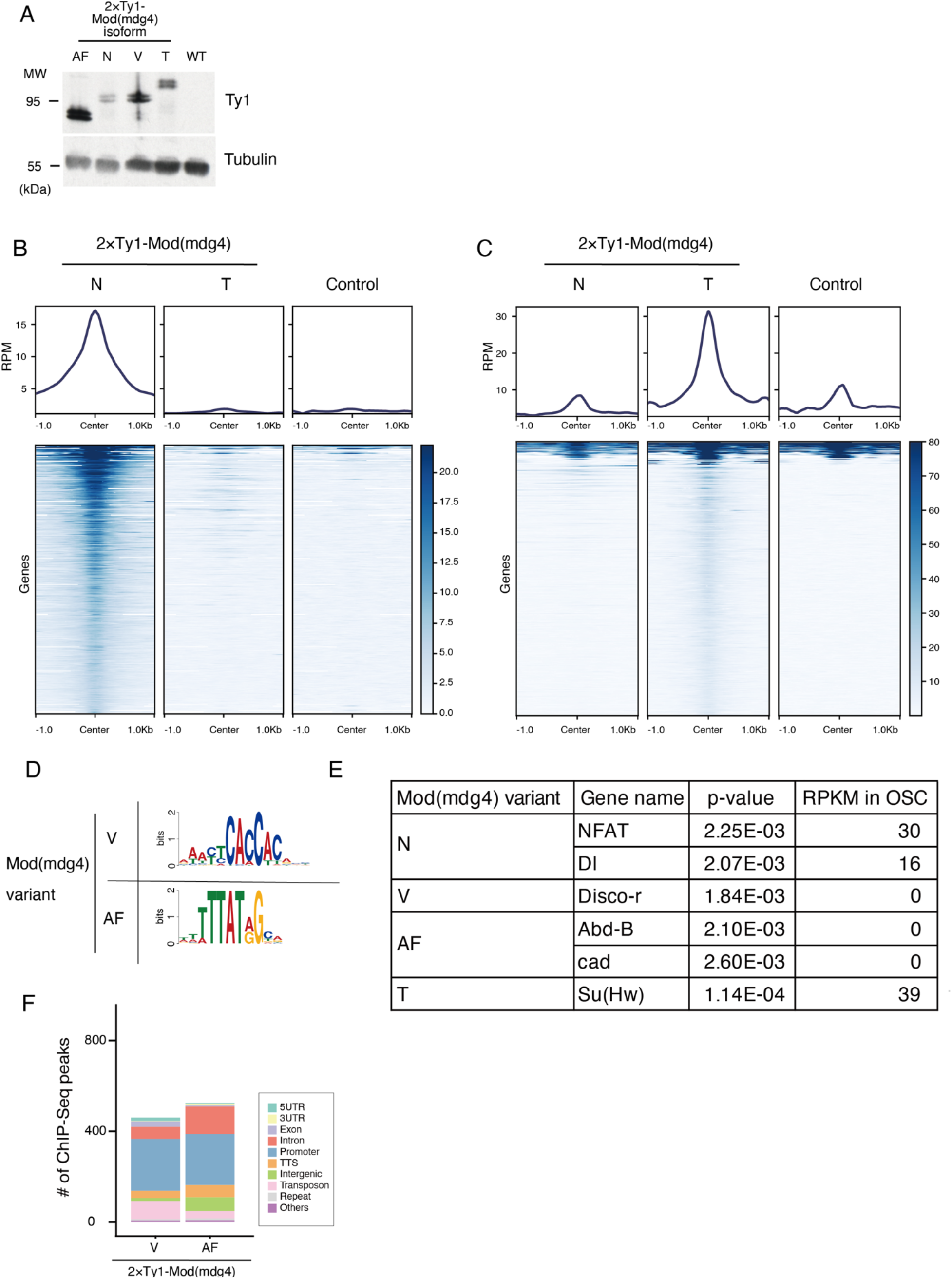
related to Fig. 2. **A**, WB by Ty1-tag antibody showing exogenous expression of 2×Ty1-tagged Mod(mdg4) AF, N, V or T in OSC. Tubulin was used for loading control. Molecular weight (MW) is indicated at the left of each image. **B**,**C**, Metaplot and heatmap indicating Mod(mdg4)-N, -T, and control (anti-Ty1 ChIP against wild-type OSC) signal level within 1.0 kb of Mod(mdg4)-N (B) or -T (C) peaks. Heatmap is sorted by intensity of Mod(mdg4)-N (B) or -T (C) signal. Y-axes of metaplots are normalized by RPM. Note that some sites of Mod(mdg4)-T peaks (C) display high background signal because Mod(mdg4)-T strongly accumulates on *gypsy* retrotransposons, which have many copies within the genome. **D**, Logos showing clear enrichment around ChIP-Seq peaks of Mod(mdg4)-V and -AF. These motifs are identified by de novo motif discovery analysis MEME (Multiple Em for Motif Elicitation). **E**, Transcription factor motifs resembling each Mod(mdg4) variant motif and its p-value identified by TOMTOM. RPKM in OSC is from modEncode RNA-Seq results. **F**, Functional annotation of peaks of Mod(mdg4)-V and -AF, identified by HOMER. The y-axis shows number of peaks of each Mod(mdg4) variant.

**Fig. S4.**
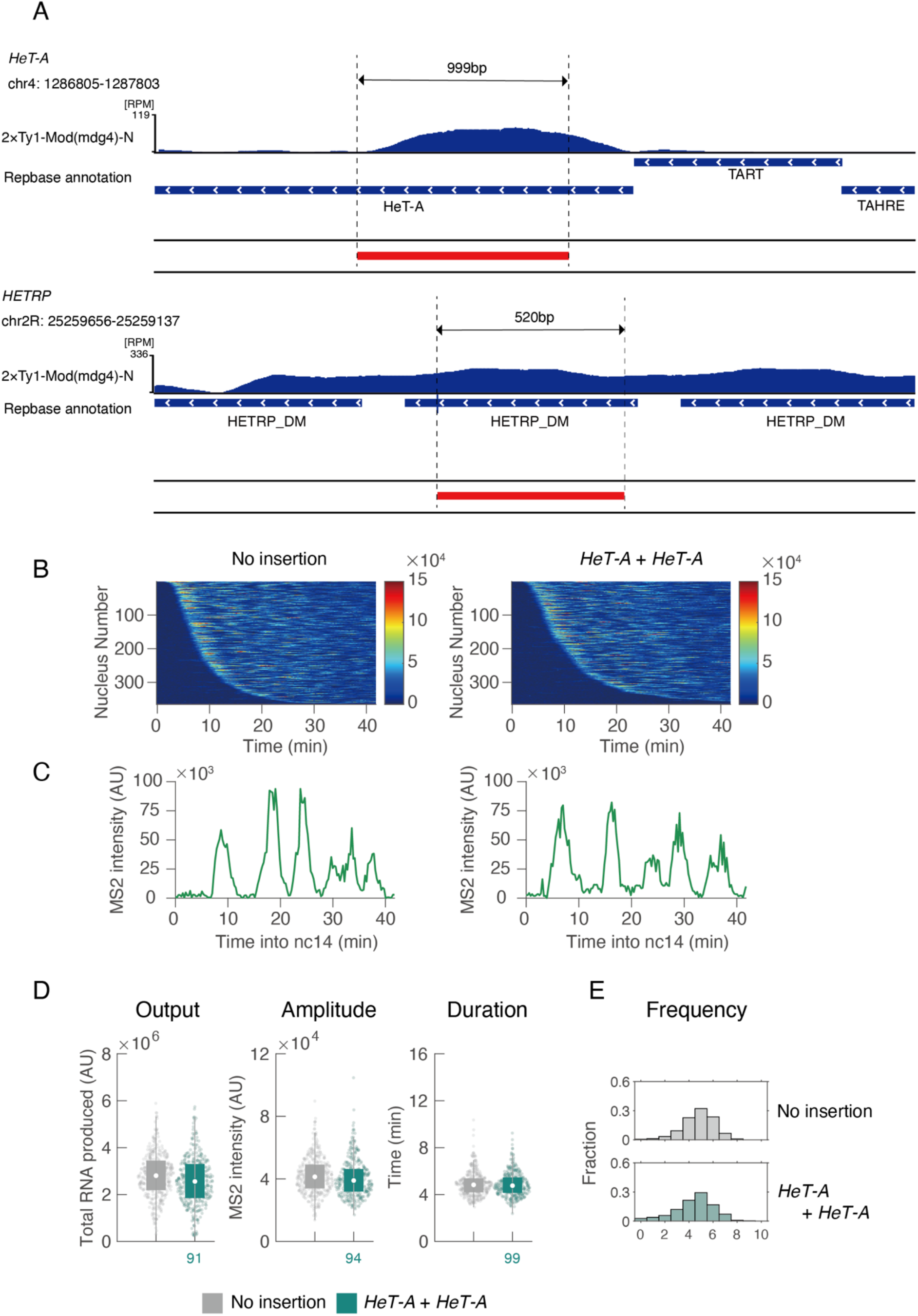
related to Fig. 3. **A**, Coverage plot showing sequences used for live-imaging of enhancer-blocking assay. chr4: 1286805-1287803 (dm6) region is referred to as *HeT-A*, and chr2R: 25259656-25259137 (dm6) is referred to as *HETRP* in this study. Red bars indicate the sequences used for enhancer-blocking assay. Mod(mdg4)-N ChIP-Seq coverage (blue) and Repeatmasker annotations of repetitive sequences are indicated. Note that there are some mutations in cloned sequences compared with reference and precise sequences used for assay are written in Supplemental Table 2. **B**, MS2 trajectories for all analyzed nuclei. Each row represents the MS2 trajectory for a single nucleus. A total of 365 and 365 ventral-most nuclei, respectively, were analyzed from three independent embryos for the reporter genes with no insertion of insulators (left) and double insertion of *HeT-A* sequence (right). Nuclei were ordered by their onset of transcription in nc14. AU; arbitrary unit. Note that plot of control is the same as the plot in Figure 3B. **C**, A representative trajectory of transcriptional activity of the MS2 reporter gene with control (left) and double insertion of *HeT-A* sequence (right). **D**, Boxplots showing the distribution of total output (left), burst amplitude (middle) and burst duration (right). The box indicates the lower (25%) and upper (75%) quantile and the open circle indicates the median. Whiskers extend to the most extreme, non-outlier data points. A total of 365 and 365 ventral-most nuclei, respectively, were analyzed from three independent embryos for the reporter genes with no insertion of insulators (left) and double insertion of *HeT-A* sequence (right). Median values relative to the control reporter are shown at the bottom. AU; arbitrary unit. Note that plot of no insertion is the same as the plot in Figure 3F. **E**, Histograms showing the distribution of burst frequency during nc14 stage. A total of 365 and 365 ventral-most nuclei, respectively, were analyzed from three independent embryos for the reporter genes with no insertion of insulators (top) and double insertion of *HeT-A* sequence (bottom). Note that plot of no insertion is the same as the plot in Figure 3G.

**Fig. S5.**
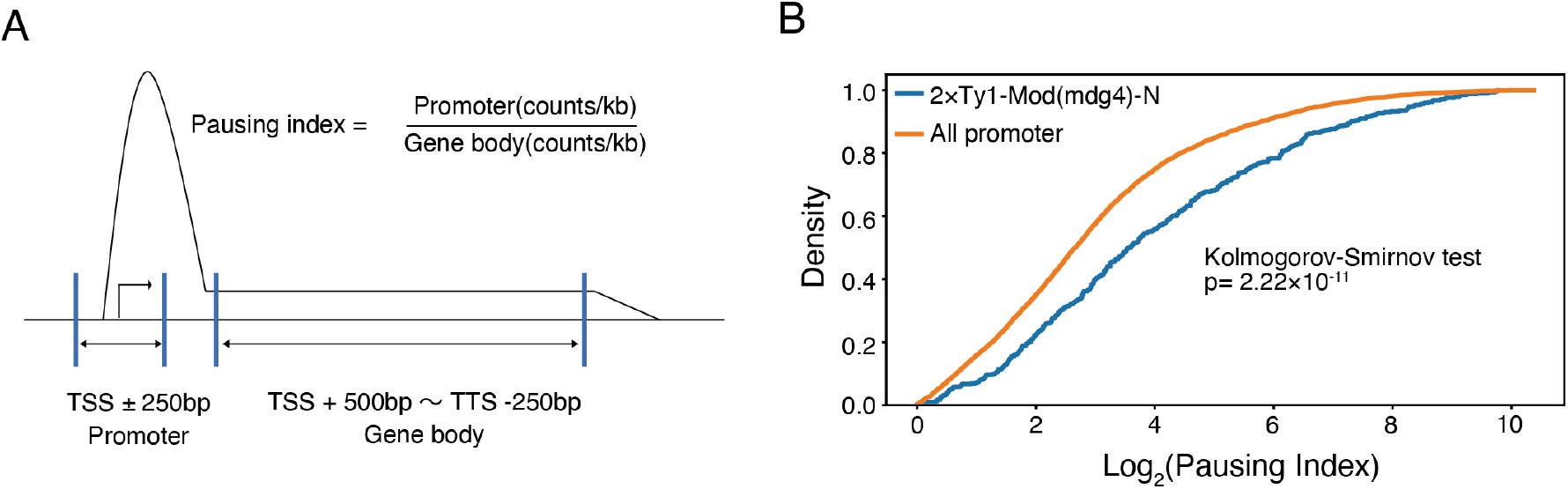
Pausing index and Mod(mdg4)-N distribution. **A**, Schematic diagram showing how pausing index is calculated. Pausing index is the value obtained by dividing the number of reads/kb in the promoter region (TSS ±250 bp) by the number of reads/kb in the gene body (TSS +500 bp ∼ TTS -250 bp) region. **B**, Cumulative distribution plot showing pausing index of all promoters (orange) and Mod(mdg4)-N bound promoters (blue) from reanalysis from Sienski G et al, 2011. (Kolmogorov-Smirnov test p= 2.22×10^−11^). The x-axis is log2 ratio.

**Fig. S6.**
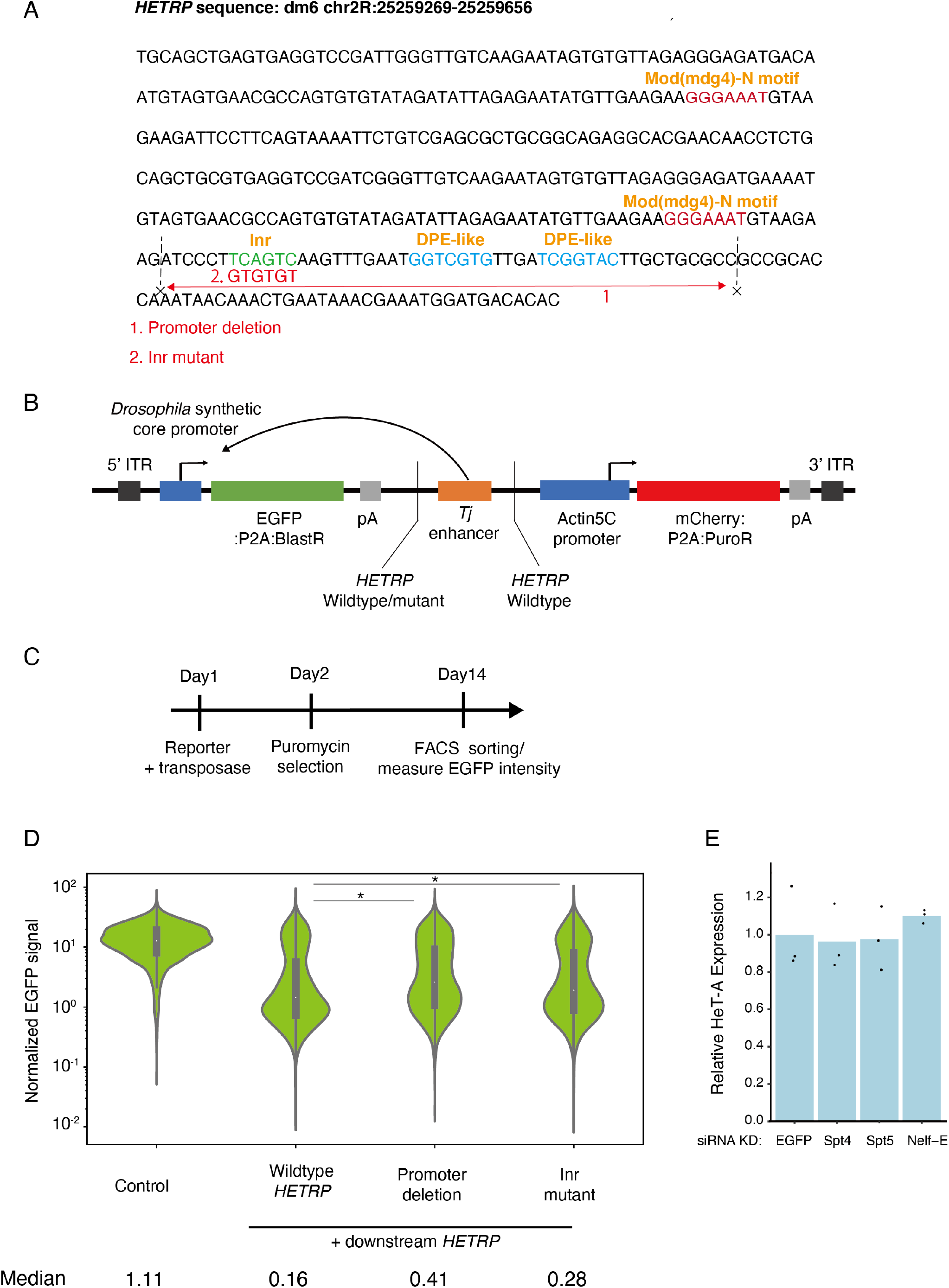
Enhancer blocking function depends on Pol II recruiting machinery. **A**, *HETRP* sequence and its mutation used for experiment. Initiator (Inr) sequence (green) and Downstream Promoter Element (DPE) sequence (blue) are indicated. Two mutants (Promoter deletion and Inr mutant) are generated from this sequence as shown (red). **B**, Schematic representation of reporter gene used for assay. Tj enhancer is located downstream of *Drosophila* synthetic core promoter-EGFP:P2A:Blastcidin-S deaminase gene. Actin5C promoter-mCherry:P2A:Puromycin N-acetyl transferase provide a selective marker. This plasmid is inserted into the genome with Piggybac transposase. **C**, Experimental design of assay. Reporter gene is integrated into the genome with Piggybac transposase. Then, successfully integrated reporter is selected with puromycin. After 13 days of selection, mCherry-positive cells are sorted and EGFP-intensity is measured with FACS. **D**, Violin plot showing the results of normalized EGFP intensity of no insertion control, downstream *HETRP* with wild-type *HETRP* insertion, promoter deletion mutant or initiator mutant, from left to right. Mean and median values of EGFP intensity are shown below. The y-axis is log10 ratio. An asterix (*) indicates the hypothesis that these two distributions are from same population is rejected at p = 0.01 significant level with Brunner-Munzel test. **E**, Bar plot showing qRT-PCR result of *HeT-A* expression upon EGFP (control), Spt4, Spt5 and Nelf-E KD in OSC.

**Fig. S7.**
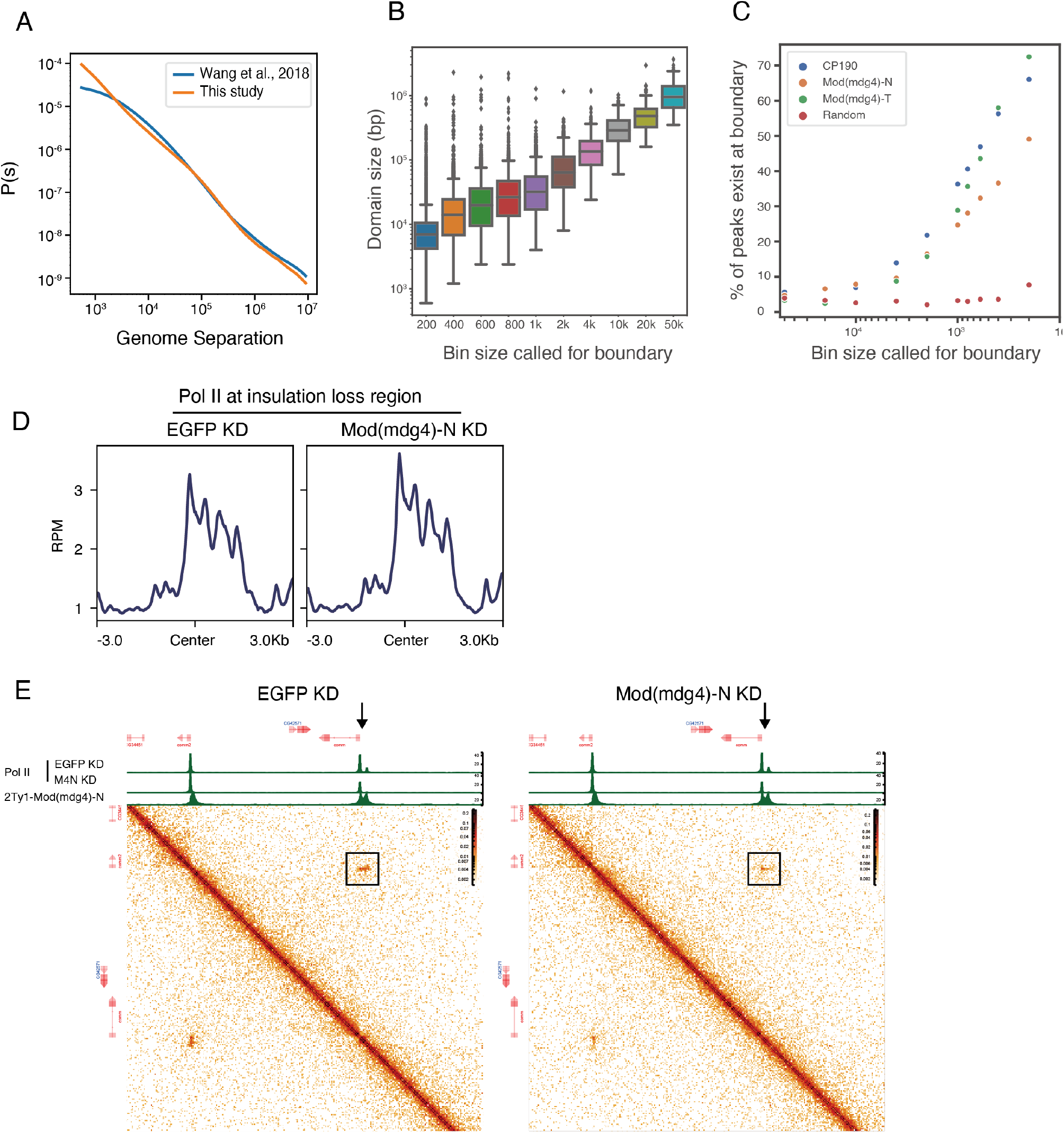
related to Fig. 5. **A**, P(s) curve comparing Wang et al., 2018 (Hi-C) and this study (Micro-C XL). **B**, Box plot showing length distribution of chromatin domain structure defined with insulation score at different resolutions (from 50 kbp to 200 bp). The y-axis is a log10 scale. **C**, Scatter plot showing proportion of ChIP-Seq peaks of CP190 (blue), Mod(mdg4)-N (orange), Mod(mdg4)-T (green) and random sequence (red) which overlap with boundaries called at each bin size. The x-axis is a log10 scale. **D**, Metaplot showing Pol II distribution around chromatin insulation lost sites over ±3.0 kb region from boundaries upon EGFP KD (left) and Mod(mdg4)-N KD (right). **E**, Another example of region where loss of Mod(mdg4)-N causes genomic looping loss. Left contact map indicates EGFP KD and right contact map indicates Mod(mdg4)-N KD. Black rectangles indicate loop between comm promoter and comm2 promoter. Coverage tracks of Mod(mdg4)-N and Pol II distribution upon EGFP (control) and Mod(mdg4)-N KD are shown at top of contact map.

**Supplemental Table 1**

Primers and siRNAs used in this study

**Supplemental Table 2**

Sequences used for reporter assays

**Supplemental Video 1**

Live imaging of MS2-yellow-sna shadow enhancer reporter with no insertion of insulators (WT), single insertion of *gypsy* insulator, *HeT-A* sequence, or *HETRP* sequence and double insertion of *HETRP* sequence from left to right during nc14. The maximum projected images of MCP-GFP and His2Av-mRFP are shown in green and red, respectively. Images are oriented with ventral view facing up.

**Supplemental Video 2**

Live imaging of MS2-yellow-sna shadow enhancer reporter with no insertion of insulators (left) and double insertion of *HeT-A* sequence (right) during nc14. The maximum projected images of MCP-GFP and His2Av-mRFP are shown in green and red, respectively. Images are oriented with ventral view facing up.

